# An Open-Source Reproducible Workflow for Pocket-Oriented Virtual Screening and ADME-Integrated Chemoinformatics: A Multi-Target Flavivirus Case Study

**DOI:** 10.64898/2026.04.28.721199

**Authors:** Juan Philippe Teixeira, Miklos Maximiliano Bajay, Caio César de Melo Freire, Lorenzo Borelli Ferraz Bettin, Anderson Pereira Soares, Daniel Ferreira de Lima Neto

**Affiliations:** State University of Santa Catarina – Center for Agricultural and Veterinary Sciences – Postgraduate Program in Biochemistry and Molecular Biology (PMBqBM); State University of Santa Catarina – CERES - Laguna; Federal University of São Carlos (UFSCar) – Department of Genetics and Evolution; São Paulo State University (UNESP); University of São Paulo (USP); Federal University of Santa Catarina

## Abstract

Zika virus (ZIKV), yellow fever virus (YFV), West Nile virus (WNV), Usutu virus (USUV), and Saint Louis encephalitis virus (SLEV) remain major public health concerns, yet broad-spectrum antiviral options are limited. Here, we present an open-source, reproducible software workflow for pocket-oriented virtual screening and ADME-integrated chemoinformatics, designed to support standardized multi-target compound prioritization. As a case study, the workflow was applied to structural and nonstructural proteins from clinically relevant flaviviruses. Automated pocket detection using Concavity reduces site-selection bias by generating docking boxes from surface concavity clusters, while standardized downstream scripts parse docking logs, convert docking-derived binding energies into Kd-related metrics, integrate SwissADME descriptors, and compute LE, LLE, FQ, and drug-likeness rules. The framework also supports retrospective validation and comparative benchmarking using literature-supported reference compounds and target-specific plausibility checks. Rather than proposing experimentally validated antiviral candidates, this study provides a reusable computational framework for hypothesis generation, benchmarking, and downstream experimental prioritization in structure-based drug discovery. The workflow is modular and adaptable to other multi-target screening campaigns where integrated ranking across binding, physicochemical, and ADME dimensions is required.

**SUMMARY:** We describe an open-source, reproducible software workflow that integrates pocket-oriented docking, ligand efficiency scoring, ADME descriptor integration, and multivariate chemoinformatics to standardize compound prioritization across multiple protein targets. The workflow combines open-source tools with auditable Bash, R, and Python scripts and is demonstrated through a multi-target flavivirus case study. Rather than claiming experimentally validated antiviral activity, the framework is intended to support hypothesis generation, retrospective benchmarking, transparent reporting, and downstream experimental prioritization.

## INTRODUCTION

This study presents a reproducible open-source computational workflow for pocket-oriented molecular docking, ADME integration, ligand-efficiency scoring, and multivariate chemoinformatics analysis. The workflow is demonstrated using structural and nonstructural proteins from five clinically relevant Flavivirus species as a case study. It is designed to standardize compound ranking and support hypothesis generation for downstream antiviral screening, rather than to provide stand-alone experimental proof of antiviral efficacy. In this workflow, we evaluate three structural proteins—Capsid (C), Pre-membrane/Membrane (prM/M), and Envelope (E)—and seven nonstructural proteins (NS1, NS2a, NS2b, NS3, NS4a, NS4b, and NS5), all recognized as priority targets for small-molecule intervention and considered a reliable foundation for subsequent docking and comparative molecular analysis1, 2. To contextualize the relevance of this workflow, we also summarize the public health significance of the included Flavivirus species and outline recent research advances that motivate the need for reproducible and scalable in silico screening approaches.

According to the International Committee on Taxonomy of Viruses, Flavivirus^3^ comprises positive-stranded, non-segmented RNA viruses (+ssRNA) with a small genome of approximately 11 kilobases and conserved molecular features. This genus includes more than 70 species capable of infecting mammals, producing a wide range of clinical outcomes from asymptomatic infections to fatal disease, including persistent post-acute manifestations and sequelae. Among these, Dengue Virus (DENV), Zika virus (ZIKV), Yellow Fever Virus (YFV), West Nile Virus (WNV), Usutu Virus (USUV), and Saint Louis Encephalitis Virus (SLEV) exhibit broad host ranges and tissue tropism closely linked to nuanced structural characteristics. Together, these traits contribute to their high adaptability and virulence, establishing them as significant human pathogens^4^.

These species have long been linked to major disease outbreaks and remain central to global epidemiological surveillance. Yet Flavivirus infections continue to strain scientific and public-health efforts due to expanding geographic ranges, resurgence patterns, underreporting, misdiagnosis, and persistent gaps in prevention and control^5^. This concern is particularly pronounced in regions where local health system constraints can lead to loss of life and substantial economic impact, underscoring the need for more efficient research tools to support the development of viable therapeutic alternatives and improve containment strategies^6^. This need is further reinforced by hurdles such as antibody-dependent enhancement (ADE), cross-reactivity, and the scarcity of molecular and *in vivo* models suitable for low-input approaches that can help translate laboratory findings into clinical solutions^7, 8^. Additionally, Flavivirus threat is expected to rise in the coming decades due to vector expansion, and other interconnected factors, emphasizing new strategies that unify drug discovery and cross-virus structural analysis to overcome current therapeutic limitations^9^.

Considering this, DENV has served as a primary model for understanding genome organization, diversity, and evolutionary mechanisms within the *Flaviviridae* family^10^. Its interactions with host factors and the immune system provide a framework for exploring viral complexity across Flaviviruses and serotypes^11^, an especially valuable asset given the absence of specific or broad-spectrum antivirals. Accordingly, the pipeline presented here is guided by DENV’s homology with the selected Flaviviruses, along with computational and experimental frameworks on phylogenetic relationships that inform pharmacological screening in this field^12^.

The first target of this workflow is ZIKV, an emerging pathogen that has prompted efforts to understand transmission, natural history, and co-evolution with human hosts13. Recently, animal models have been developed to test vaccine candidates and non-antibody-dependent strategies, often complemented by computational and mathematical tools that predict viral spread based on molecular analyses14–16. ZIKV shares substantial genomic, structural, and immunological similarities with DENV; thus, understanding ZIKV molecular mechanisms in comparison with other Flaviviruses is essential for improving antiviral strategies. Here, binding modes of screened natural compounds in ZIKV proteins were compared to their homologous counterparts in other Flaviviruses, revealing consistent trends in predicted antiviral activity.

YFV, the next target, has caused devastating epidemics since the seventeenth century and remains a major threat in Africa and South America17. Although an effective live-attenuated vaccine has been available since the 1930s18, the unprecedented 2016–2019 epidemic in Brazil demonstrated the virus’s capacity to re-emerge in areas previously considered YF-free, prompting intensified genomic monitoring19. As a result, recent research has shifted from traditional epidemiological surveillance and vector control to real-time sequencing supported by large-scale data integration20. However, limited vaccination coverage and YFV’s marked hepatotropism underscore the need for non-hepatotoxic antiviral alternatives. These can be comprehensively explored using the natural-compound screening approaches implemented in this workflow, which also build upon previous evidence showing the antiviral activity of compounds such as sofosbuvir21.

Also included in this pipeline, WNV is a globally distributed Flavivirus responsible for substantial human and animal morbidity. Its epidemiological profile changed in the mid-1990s following major outbreaks and the subsequent expansion across the Western Hemisphere. Despite extensive research, no licensed vaccines or targeted antiviral treatments are available, and patient management remains supportive^22^. Knowledge gaps persist regarding viral distribution, genetic variability, and host–vector interactions, many of which can be computationally explored through integrated molecular docking and structural modeling by expanding insights from previous, more restricted screenings^23, 24^ and immunoinformatics^25^. In addition, the provided protocol incorporates statistical inference for pharmacological profiling, supporting subsequent investigations that are consistent with robust molecular dynamics methodologies^26^.

USUV, the fourth target, is an emerging Flavivirus that has gained attention due to its expanding range and zoonotic potential in recent decades^27^, prompting increased interest in viral evolution and phylogenetics^28^. Recent molecular studies have identified distinct viral lineages involved in dispersal and maintenance^29^; however, implementation of a true “One Health” framework for monitoring and control remains limited. No approved or widely validated therapeutic options currently exist for USUV infections, emphasizing the need for strengthened surveillance and comparative molecular studies^30^.

The last considered target is SLEV, a neurotropic Flavivirus that likely originated in Central America and remains a major cause of viral encephalitis in the region^31^. Additional foci occur in the southwestern United States, with notable re-emergence and outbreaks reported in Argentina (2005), São Paulo (2006), and Arizona (2015)^32–34^. Currently, no licensed vaccines or specific antiviral treatments are available for SLEV, and clinical management remains supportive, although preliminary studies have identified promising antiviral candidates^35^. Recent computational advances in multi-epitope vaccine design are encouraging^36^, reinforcing the value of cross-viral analyses for elucidating shared pathogenic pathways and pharmacological interactions for drug development guidance.

Building on these insights, the proposed in silico cross-docking protocol integrates natural compounds with homologous proteins from ZIKV, YFV, WNV, USUV, and SLEV, offering broader target coverage and epidemiological relevance over single-target approaches. The rationale emphasizes natural compounds as valuable bioactive sources with enhanced chemical diversity, complexity, and biocompatibility, supporting unbiased exploration of chemical space37, while synthetic drugs were also docked and evaluated for comparison (e.g., sofosbuvir). This systematic, standardized workflow unifies compatible open-source tools within a pragmatic framework, addressing common limitations of ad hoc analyses with inconsistent metrics and fostering collaborations in computer-aided drug discovery (CADD)38. The free software usage reinforces the workflow’s cost-effectiveness and enables rapid screening of large compound libraries across multiple viral targets, reducing reliance on early in vitro and in vivo testing, which in turn extends current directions in Flavivirus research to support discoveries on neglected and emerging species39, 40. This computational methodology also combines ligand-based and structure-based techniques that could help anticipate resistance mutations in structural molecules, following recent trends regarding other viral systems41, 42. This approach also enables the identification of multi-variable interaction fingerprints for machine learning applications 43.

Moreover, this fully auditable pipeline automates pocket-specific docking box generation using Concavity-based detection of surface depressions from residue centroids^44^, reducing common site-specific bias. The R-based multidimensional analyses (heatmaps, ΔG clustering, 3D scatter plots), ΔG-to-Kd thermodynamic conversion, and pharmacokinetic filtering using curated ADME datasets and ligand efficiency metrics (LE, LLE, FQ) also enhance reproducibility compared to traditional static-grid docking^45, 46^. The built-in multivariate analysis module (PCA, t-SNE, UMAP, LDA) further integrates binding, structural, and ADME-Tox parameters for compound ranking based on multidimensional profiles, aligning with established best practice in multi-omics and chemometrics to address analytical fragmentation^47^. Designed for efficiency, this framework minimizes manual input while allowing direct integration with advanced macromolecular modeling and QSAR virology workflows^48–51^.

In this article, we describe a reusable software workflow that provides Bash and R scripts with detailed documentation to enable both new and experienced users to conduct multi-target molecular docking, ADME integration, ligand-efficiency scoring, and compound prioritization analyses with customizable functions and commands. All computational tools are open-source and freely accessible. The complete workflow, schematically represented in Figure 1, includes automated binding pocket detection, molecular docking, thermodynamic conversion, ligand efficiency metrics calculation, ADME property integration, drug-likeness rule evaluation, and multivariate dimensionality reduction analyses for compound ranking across multiple viral protein targets. This workflow enables cross-referencing results against established ligand efficiency benchmarks and ADME-Tox descriptors, serving as a practical software framework for computational chemistry practitioners and wet lab researchers implementing virtual screening workflows for multi-target drug discovery.

**Figure 1:**
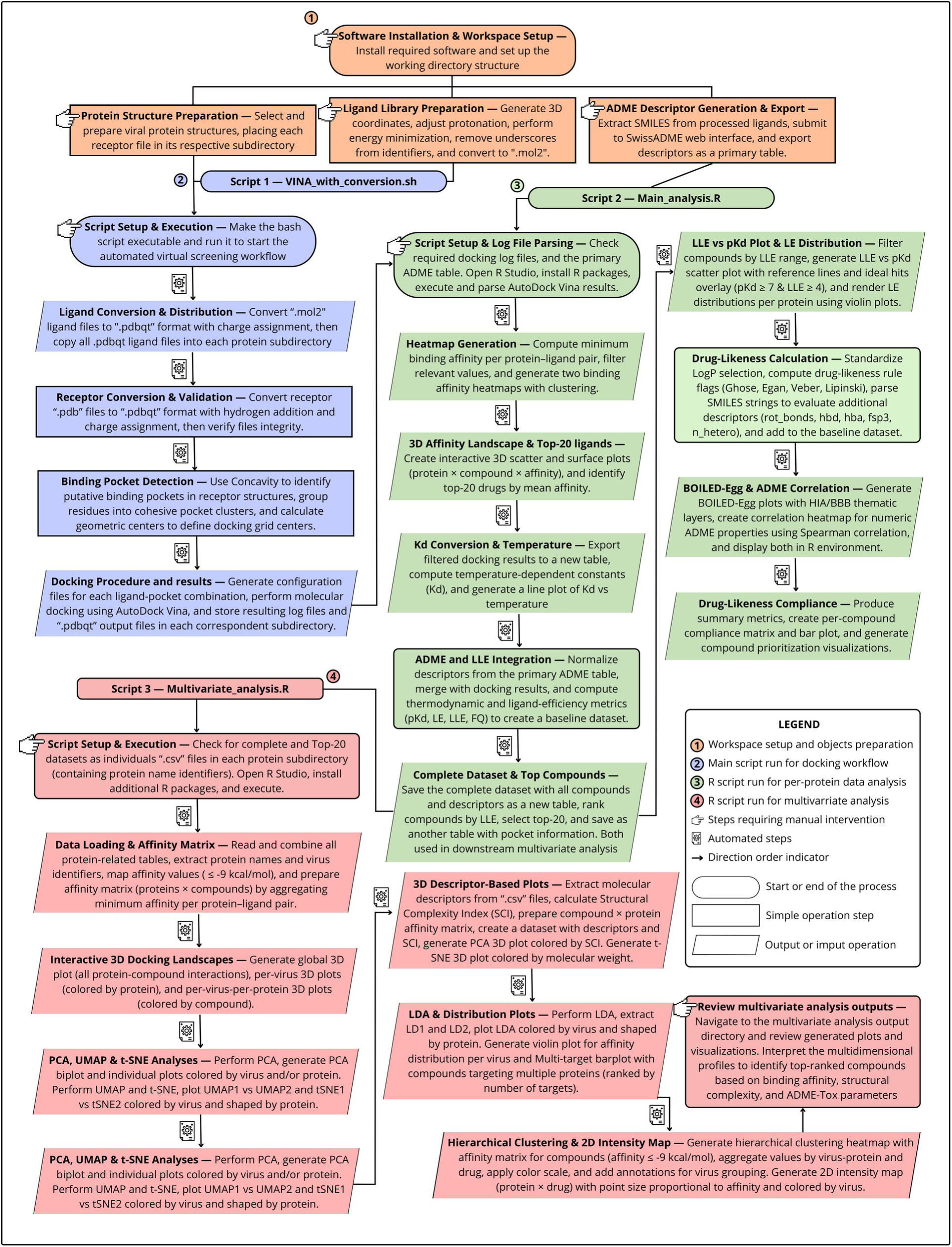
Schematic summary of the Automated Pocket-Oriented Docking and ADME-Integrated Analysis for Multi-Target Flavivirus Screening pipeline. This flowchart is divided into four major sections: (1) workspace setup and object preparation (orange), (2) automated docking workflow execution (blue), (3) per-protein data analysis and ligand efficiency scoring (green), and (4) multivariate analysis and visualization (red). Each step shows the corresponding section number and a brief description. Plain arrows indicate direct flows between steps. Steps requiring manual intervention are indicated according to the figure legend. This pipeline provides automated molecular docking with binding pocket detection, comprehensive ligand efficiency metrics (LLE, LE, FQ), ADME property integration, drug-likeness rule evaluation, and multivariate dimensionality reduction analyses (PCA, UMAP, t-SNE, LDA) for compound prioritization across multiple viral protein targets. The workflow is implemented as bash and R scripts in a step-by-step manner, enabling users to learn and practice virtual screening and ligand efficiency analysis.

## METHODS

NOTE: Information related to the software applications, versions, and dependencies used in this study is listed in the **Table of Materials**. Troubleshooting guidance for common issues encountered during protocol execution is provided in the Discussion section (Table 5).

### 1. Downloading and Installing Necessary Software and Utilities

NOTE: This workflow is designed and tested for Linux systems. The bash scripts, command-line tools (Antechamber, OpenBabel, Concavity), and path conventions (e.g., ∼/Vina_Workspace) are optimized for Unix-like environments. For Windows users, running this workflow requires Windows Subsystem for Linux (WSL), which provides a Linux-compatible environment. Adapting to other operating systems may require rewriting shell scripts, modifying file path handling, and ensuring equivalent tool availability, which is beyond the scope of this workflow.

1.1. Download AutoDock Vina^52^ from: https://github.com/ccsb-scripps/AutoDock-Vina/releases. Follow the installer instructions provided on the GitHub release page.

1.2. Download AutoDock4^53^ from: https://autodock.scripps.edu/download-autodock4/. Follow the installation instructions on the official site.

1.3. Download OpenBabel^54^ from: https://openbabel.org/index.html. Follow the installation guide to install the latest version.

1.4. Download AmberTools^55^ package from: https://ambermd.org/GetAmber.php#ambertools. Follow the installation guide to install the latest version.

1.5. Download Python^56^ from: https://www.python.org/downloads/. Install the latest stable release, ensuring to add Python to your system PATH.

1.6. Download R binaries and RStudio^57^. R: https://cran.r-project.org/. RStudio: https://posit.co/download/rstudio-desktop/. Install R first, then RStudio, following the respective installation instructions.

NOTE: R Packages will be installed and loaded automatically via the R script provided in the workflow. Ensure R is properly configured to install packages (from CRAN, Bioconductor, etc…) as needed.

1.7. Download Pymol from https://www.pymol.org/. Follow the manual installation instructions provided on the website. Optionally download and install APBS Algorithms from: https://www.poissonboltzmann.org/.

NOTE: This APBS tool can be used for manual curation and is compatible with the molecular visualization program Pymol^58^ as an add-on plugin^59^.

1.8. Download all the files used in this pipeline. Access the GitHub repository: https://github.com/Thermodynamik-und-Strukturkomplexe/Vina-SBVS-LBVS-ADME-Full-Workflow-Documentation. Download the entire repository as a ZIP file or clone it using Git.

1.8.1. To download the ZIP file from the repository in Unix-like systems, use the following command in Bash: **wget** https://github.com/Thermodynamik-und-Strukturkomplexe/Vina-SBVS-LBVS-ADME-Full-Workflow-Documentation/archive/refs/heads/main.zip **-O ∼/Desktop/Vina-SBVS-LBVS_ADME.zip**

1.9. Locate the Scripts inside the Vina-SBVS-LBVS_ADME folder on your Desktop. In Linux, navigate to: **cd ∼/Desktop/Vina-SBVS-LBVS_ADME/Vina_Scripts**. Ensure this folder contains “VINA_with_conversion.sh” and two R scripts for data analysis: “Main_analysis.R” and “Multivariate_analysis.R” that will be used later. If you can’t find these files, check your internet connection and download the repository again.

1.10. Create a working directory, setting up folders for receptors (proteins) and compounds: **mkdir ∼/Vina_Workspace**. Inside, create subfolders: **mkdir ∼/Vina_Workspace/receptors** and **mkdir ∼/Vina_Workspace/compounds**.

CRITICAL STEP: Adapt the Vina script for your setup. Open “VINA_with_conversion.sh” in a text editor or similar program (e.g., Visual Studio Code). Locate and update the following path variable: MGLTOOLS_UTILS: Replace **/home/your_path/…/MGLToolsPckgs/AutoDockTools/Utilities24** with the actual path to your MGLTools AutoDockTools Utilities directory on your system.

NOTE: The script uses hardcoded paths for the compounds’ and receptors’ directories. If you follow the standard directory structure, no manual configuration of these paths is required. Modifying these paths is not recommended and should be attempted only by advanced users who understand the implications for the workflow.

### 2. Viral Protein Structure Preparation

NOTE: This workflow assumes users already have available and reliable protein models to use as primary inputs. Several computational tools can generate reliable 3D protein models from primary sequence data to ensure model quality and reproducibility60–63. These tools may be used individually or combined according to user preference, provided all used models are in “.pdb” format and without any structural/functional relevant gap.

2.1. Select viral structures using available crystallographic data or high-confidence homology models for all structural (C, prM/M, and E) and non-structural proteins (NS1, NS2A, NS2B, NS3, NS4A, NS4B, and NS5) from ZIKV, USUV, YFV, SLEV, and WNV.

2.3. Place a single protein structure in “.pdb” format named “receptor_{PROTEIN_NAME}.pdb” inside each protein subdirectory. Edit {PROTEIN_NAME} to match the subdirectory name (e.g., **∼/Vina_Workspace/receptors/WNV_E/receptor_WNV_E.pdb**).

NOTE: Each protein subdirectory must contain its own receptor file that matches this naming convention, which prevents errors from accidentally using the wrong receptor structure for different protein targets in the automated script runs.

### 3. Ligand Library Composition and primary ADME table export

3.1. Assemble a focused library, combining natural molecules and approved or investigational drugs that possess known or potential antiviral activity.

NOTE: This study employed approximately 130 compounds (Supplementary material, file 4) that were filtered during the automated screening processes. Ligands should be obtained from external 3D databases (e.g., PubChem^64^) and manually curated beforehand to remove structures with evidently poor pharmacokinetic properties^65^. After the library preparation, the screening process is compatible with the descriptors produced by SwissADME^66^ generated from “.sdf” files. Any other external tool capable of exporting ADME descriptors from different file formats to “.csv” may be used, provided conventions for ADME properties remain consistent for later steps.

3.1.1. Download the ligand structures. Place all “.sdf” files in the local Downloads folder (or equivalent) and copy them to the working directory using: **cp ∼/Downloads/*.sdf ∼/Vina_Workspace/compounds/**

**CRITICAL STEP**: Identify and convert any ligand lacking valid 3D coordinates before protonation adjustment or energy minimization. Ensure such original files have a traceable prefix (e.g., “2D_{ligand_name}.sdf”) and then generate 3D geometries only for the files beginning with this prefix by using the following commands:

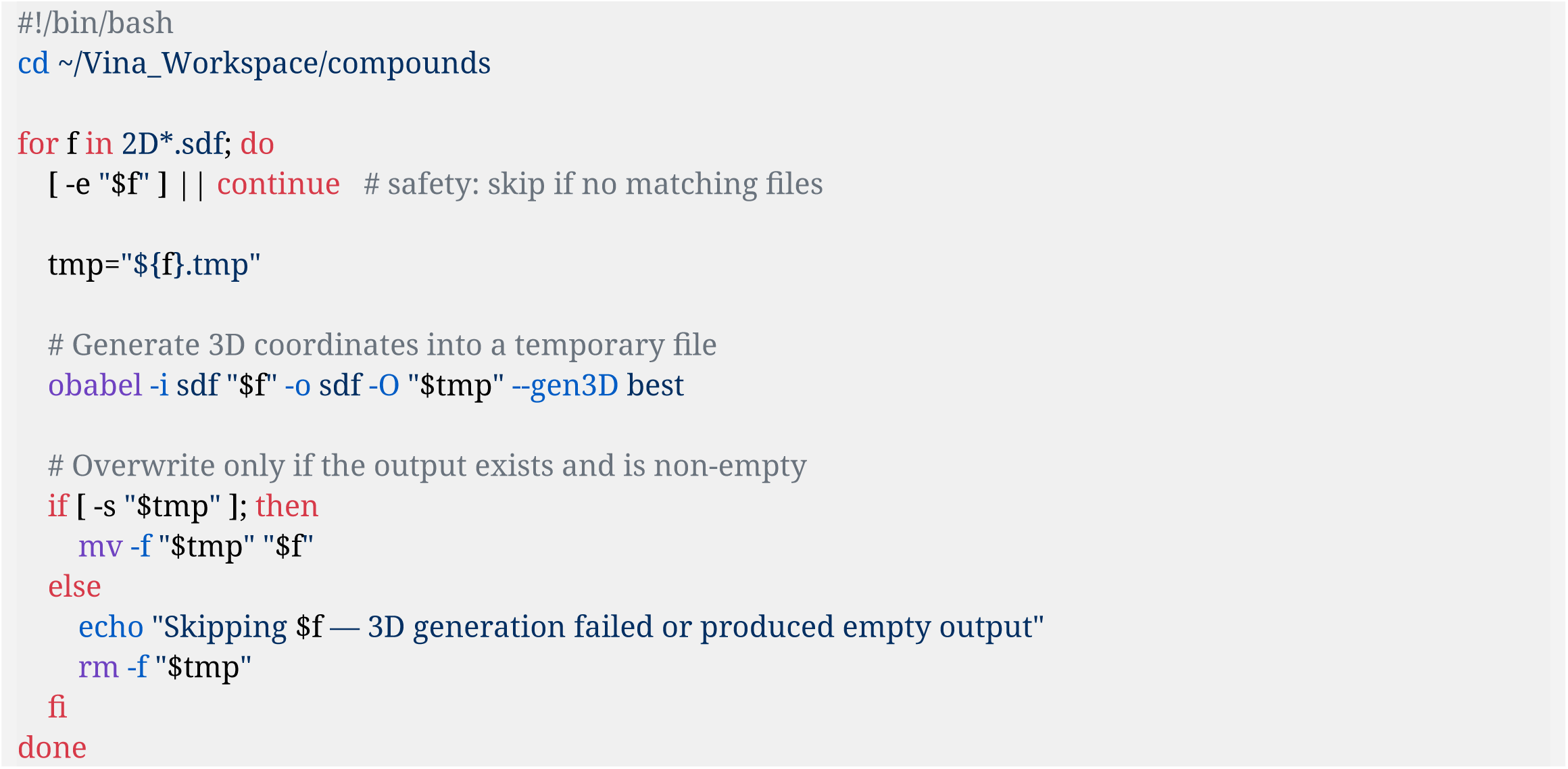

3.1.2. Run the following command in the directory containing the “.sdf” files to generate pH-adjusted structures at pH 7 (physiological):

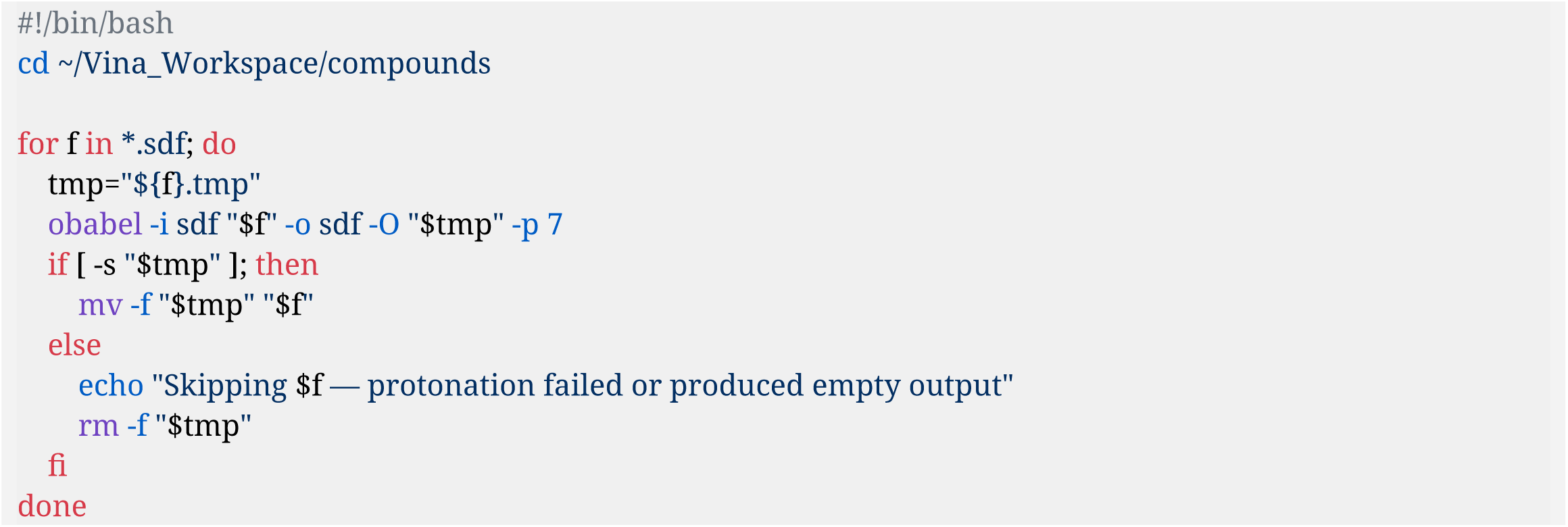

NOTE: Energy minimization relies on a force-field evaluation of all atoms and bonds ^67^. Incorrect protonation before this step can lead to the formation of an artificial, nonphysical minimum, thereby compromising the interpretation of subsequent results.

3.1.3. Run the following command in the directory containing the SDF files to generate 1,000 conformers. Select the lowest-energy weighted conformer, and minimize it for 1,000 steps:

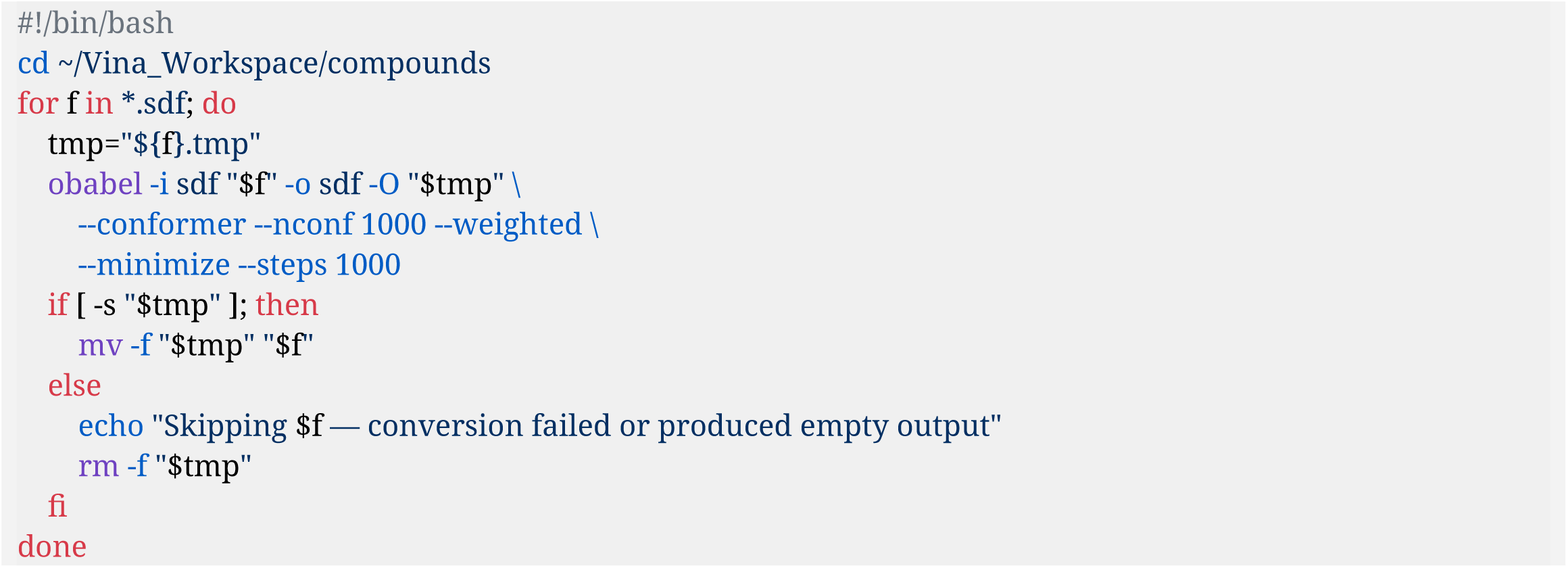

NOTE: This bash loop writes each processed structure to a temporary file before replacing the original, ensuring data integrity. If Open Babel fails or produces an empty or corrupted output, the original “.sdf” file remains untouched. With these commands, only valid, non-empty output files are used to overwrite the originals. After this minimization step, the user can evaluate the final energy of each compound using Open Babel’s **--energy flag**. Each ligand energy can be assessed using the following command (consider the “ligand_name” as a placeholder, e.g., “Temoporfin.sdf”): **obabel ligand_name.sdf -otxt --energy --append “Energy“**.

CRITICAL STEP: Check the compounds directory and remove underscores from ligand identifiers (e.g., “COMPOUND_CID_60751.sdf” to “COMPOUNDCID60751.sdf”). Convert “.sdf” ligand structures using Antechamber (part of the AmberTools^55^ package) in a loop:

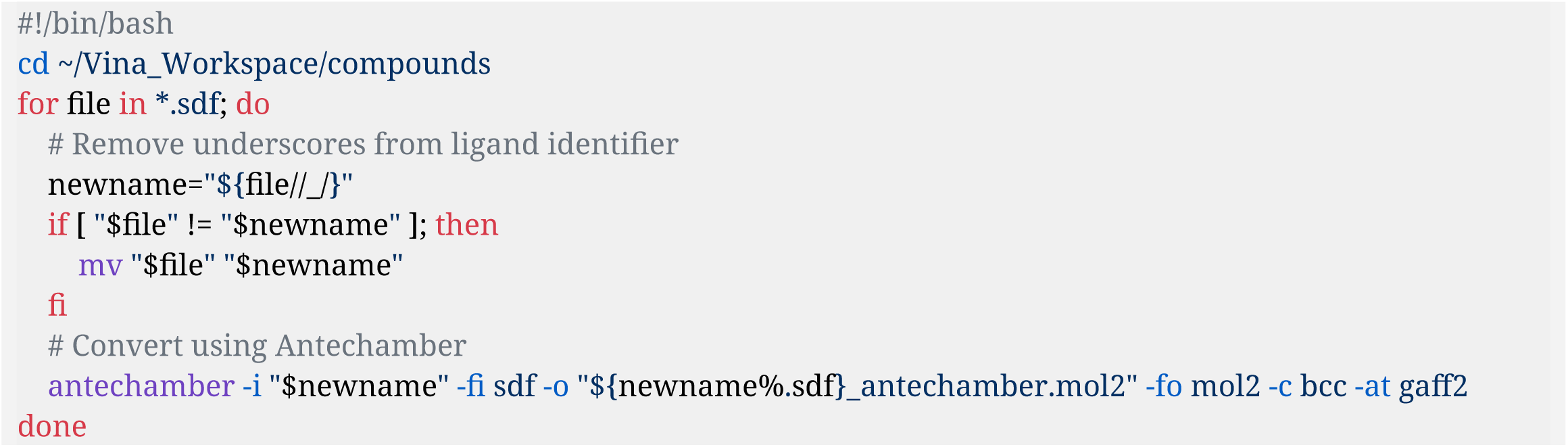

NOTE: The R analysis script (“Main_analysis.R”) used in section 5 parses log filenames by splitting on underscores. If ligand identifiers contain underscores, the parsing will incorrectly separate the protein and ligand names.

3.2. Generate “adme_input.smiles” with one SMILES per ligand (*_antechamber.mol2) using Open Babel: **obabel *_antechamber.mol2 -O adme_input.smiles -osmi –title**.

NOTE: The retained ligand identifiers in the “.smiles” file are as originally assigned in the database (e.g., ID: 60751 for Temoporfin). Therefore, users are advised to create and save a separate reference table containing a customized ID for traceability before proceeding. An example table containing standard identifiers is inside the Supplementary Material (File 3) as a spreadsheet “Compound_codes” in “.xlsx” format.

3.3. Open “adme_input.smiles” in a text editor, copy all contents, and paste into the SwissADME web interface (https://www.swissadme.ch/) in the right blank space designated for SMILES text input. Run the prediction and wait for the ADME results to be generated.

3.4. Export the SwissADME results as CSV (Download → “Export table as .csv”), and rename the file to “adme_features_for_efficiency.csv”. Copy the exported “adme_features_for_efficiency.csv” into the compounds’ directory using this command in the bash shell: **cp ∼/Downloads/adme_features_for_efficiency.csv ∼/Vina_Workspace/compounds/**.

NOTE: The exported table must contain columns with drug name, SMILES, and ADME descriptors (molecular weight, LogP variants, etc.). An example table with the complete set is provided in the Supplementary Material (File 4).

### 4. Setting up the bash Script run

NOTE: Once all input files are prepared, the provided Bash script (“VINA_with_conversion.sh”) automatically executes the complete virtual screening workflow by sequentially iterating through each receptor subdirectory. This automated pipeline includes MGLTools-based receptor and ligand preprocessing, concavity-based binding pocket detection, and molecular docking using AutoDock Vina. This full script is provided in the Supplementary Material (script 1) and is compatible with Unix-like operating systems. From this point onward, any step executed automatically by a script (regardless of the scripting language) is explicitly labeled as “(Automated)” throughout the workflow to ensure procedural clarity and reproducibility.

4.1. After downloading the VINA_with_conversion.sh bash script, make the script executable in the bash shell: **chmod +x ∼/Desktop/Vina-SBVS-LBVS_ADME/Vina_Scripts/VINA_with_conversion.sh**.

CRITICAL STEP: Before running the script, verify that the environment has sufficient read/write permissions and that no aliases or functions from **.bashrc** interfere with execution. Ensure all required tools (Concavity, AutoDockTools, and AutoDock Vina) are installed and accessible before proceeding.

4.2. To execute, run the script using its full path: **∼/Desktop/Vina-SBVS-LBVS_ADME/Vina_Scripts/VINA_with_conversion.sh**.

NOTE: The script can be run from any location. It will automatically locate and navigate to **∼/Vina_Workspace/receptors/** and **∼/Vina_Workspace/compounds/**, and convert all *_antechamber.mol2 ligand files to *.pdbqt format in the compounds’ directory (once, before processing any proteins).

### 4.3. Receptor and Ligand files conversion with charges computation

4.3.1. (Automated): Convert all ligand files (“*_antechamber.mol2”) to “.pdbqt” format in the compounds’ directory using **prepare_ligand4.py**. Assign charges to all ligand atoms in the process (typically Gasteiger by AutoDock Tools default). Skip conversion if the “.pdbqt” file is newer than the source “.mol2” file.

4.3.2. (Automated): Copy all ligand files (“*.pdbqt”) from the compounds’ directory into each protein directory using: **cp “$COMPOUNDS_SRC“/*.pdbqt**.

NOTE: After section 4.4, the script loops through each “.pdbqt” ligand file, processes it separately, and creates individual log files for each compound as “{protein}_{ligand}_pocket{idx}.log”. This allows each ligand to be docked separately with ligand-specific results (e.g., “WNV_E_Teriflunomide_pocket0.log”).

4.3.3. (Automated): Convert “receptor_{PROTEIN_NAME}.pdb” files to “.pdbqt” format using **prepare_receptor4.py**, by adding hydrogens and assigning charges (typically Kollman by AutoDock Tools default). Maintain original “.pdb” receptor files and create new “pdbqt” receptor files in each protein subdirectory with their respective names embedded (e.g., “receptor_WNV_E.pdbqt” in the WNV_E subdirectory).

4.3.4. (Automated): Verify file integrity and charge assignment for both receptors and ligands. For receptors, count atoms using **grep -c “×ATOM” receptor_{PROTEIN_NAME}.pdbqt.** For ligands, verify atom count and charge presence (detecting patterns with [+-][0-9]).

NOTE: All warnings and errors regarding “.pdbqt” conversions are automatically logged to “Preparation_warnings.txt” in **∼/Vina_Workspace/compounds/Preparation_warnings.txt**.

CRITICAL STEP: During the script run, confirm that the output “.pdbqt” files are being generated. Ensure that receptor “.pdbqt” files contain complete Cα atom coordinates and consistent residue numbering by visualizing them in Pymol. Missing or misaligned numbering can lead to docking errors and mismatched pocket definitions during subsequent analyses. Check the “Preparation_warnings.txt” file after script completion to identify any conversion or verification issues.

### 4.4. Concavity pocket recognition and Docking procedure

4.4.1. (Automated): Use the cavity detection algorithm Concavity to locate putative binding pockets in the receptor structures: **concavity receptor_{PROTEIN_NAME}.pdb concavity_output.txt.scores**.

NOTE: Concavity requires the original “.pdb” file format. The “.pdbqt” version is required for AutoDock Vina. Pocket center calculation (step 4.4.2) reads Cα coordinates directly from the “.pdb” file using a format-specific parser.

4.4.2. (Automated): Extract residues forming high-scoring concave regions from the output file “concavity_output.txt.scores”.

NOTE: Residues are grouped into cohesive pocket clusters based on spatial proximity (threshold = 0.1), without manual curation. For each detected pocket, the script calculates the geometric center by averaging the Cα coordinates from the receptor. This centroid defines the docking grid center for AutoDock Vina.

4.4.3. (Automated): Generate a dedicated configuration file for each ligand-pocket combination “config_{ligand}_pocket_{n}.txt”, containing grid coordinates and docking parameters (an example “.txt” is in the Supplementary Material, file 2). Perform docking using AutoDock Vina syntax and algorithms.

NOTE: The “VINA_with_conversion.sh” updates a master report (“Final_report.txt”) with status, coordinates, and residue clusters in the parent receptor folder **∼/Vina_Workspace/receptors/**. This report enables easy tracking of all docking events.

PAUSE POINT: At this stage, wait for the Docking runs targeting each protein. Respective resulting logs and “.pdbqt” files are generated and stored in the “**dockings/**” directory within each protein subdirectory. After each iteration, reviewing automated cavity-detection results in Pymol is advised to confirm grid accuracy and biological relevance. Electrostatic surface calculations using the Adaptive Poisson-Boltzmann Solver (APBS)^68^ are also recommended to ensure that key functional regions, especially the catalytic sites of NS3 and NS5, are fully encompassed by the docking grids. Additionally, basic knowledge of protein structure terminology is required. Consulting relevant scientific literature will aid in understanding and comparing protein-specific structural features.

4.5. Retrospective validation and benchmarking (recommended for method-focused submissions) NOTE: The workflow is designed for hypothesis generation and comparative profiling. For method-oriented journals, we recommend including a retrospective validation step that demonstrates ranking enrichment of literature-supported reference compounds and the robustness of pocket detection.

4.5.1. Assemble a small reference set (e.g., 10–30 compounds) with published activity or prior in vitro/in vivo evidence against flaviviral targets (e.g., NS3 and NS5 inhibitors, entry inhibitors). Keep these compounds flagged in the library.

4.5.2. Run the complete workflow unchanged. Quantify enrichment by reporting whether reference compounds appear among the top-N ranked ligands per target and by summarizing rank-based metrics (e.g., recall@N or enrichment factor) across key proteins.

4.5.3. Perform a baseline comparison on a subset of targets by docking the same library using a conventional fixed-grid strategy (manual or canonical site definition) and compare rank stability and reference-compound enrichment to the pocket-oriented approach.

4.5.4. Report plausibility checks for representative complexes (e.g., NS5 polymerase site) by visual inspection of poses and electrostatic complementarity, and discuss failure modes (e.g., highly lipophilic binders with low LLE).

### 5. R-Based Docking Data Analysis, Binding affinity, and Heatmap generation

NOTE: At this stage, the automated R script “Main_analysis.R” will be applied for data analysis. After docking, the script parses AutoDock Vina log files to extract binding affinities, computes minimum ΔG values per protein–ligand pair, and generates interaction heatmaps. This R script is provided in the Supplementary Material (script 2).

5.1. Check each docking subdirectory (e.g., **∼/Vina_Workspace/receptors/{PROTEIN_NAME}/dockings/**) contains all log files (e.g., “{PROTEIN_NAME}_{LIGAND_NAME}_pocket{idx}.log”). Check if “Final_report.txt” is located at **∼/Vina_Workspace/receptors/Final_report.txt** (the main docking report, created at the receptors level). Check if “adme_features_for_efficiency.csv” (prepared in step 3.5.3) is in **∼/Vina_Workspace/compounds/**.

CRITICAL STEP: Open RStudio. Go to File -> Open File. Find “Main_analysis.R” inside **/Desktop/Vina-SBVS-LBVS_ADME/Vina_Scripts** and load it. Install required R packages if not already installed by running the following command in the R console:

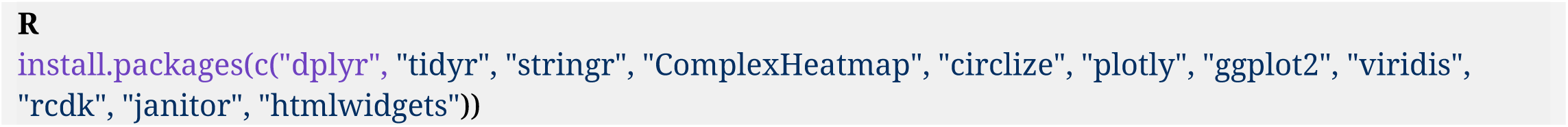

NOTE: The “Main_analysis.R” script detects and processes all protein directories with dockings/ folders containing “.log” files. The script processes each protein sequentially in a loop (e.g., ∼/Vina_Workspace/receptors/WNV_E/), producing separate outputs (graphs and other files) for each target in their respective directories. No manual setwd() or protein naming configuration is required if the user followed the standard conventions in this workflow.

5.2. Execute the entire script in the R console. Select all text inside “Main_analysis.R” script with Ctrl+A and press Ctrl+Enter (or click “Run”).

NOTE: As an alternative to executing the entire script at once, individual analysis blocks within the script may be run separately by selecting the desired code section and pressing Ctrl+Enter in the R console. This approach is recommended for users who are not yet familiar with the workflow or who wish to inspect, modify, or debug specific code blocks.

5.3. (Automated) Load required R packages (e.g., **stringr** and **dplyr**). Set working directory based on **VINA_WORKSPACE** variable. Retrieve all “.log” files from the **dockings/** subfolder using **list.files()**.

5.4. (Automated) Define a custom function that reads a single log file, extracts the protein, ligand, and pocket identifiers, and isolates the minimum binding affinity (kcal/mol) from the first docking mode. Return results as a **tibble()**. Apply this function to all log files to create **df_results**.

5.5. (Automated) Load additional R packages (e.g., **ComplexHeatmap**). Compute the minimum binding affinity per protein–ligand pair to create **df_min**, preserving the pocket information corresponding to the best binding affinity. Filter the results to retain plausible values with **filter(min_affinity > −20 & min_affinity < 0)** to create **df_filtered**.

5.6. (Automated) Reshape the data into a matrix format, converting it to a numeric matrix (**mat_filt**). Create a color scale and generate two heatmaps **(ht1 and ht2)** with **Heatmap()**: one with protein as row and drugs as columns, and an “inverted” version with drugs as rows and protein as column. Add overall titles, render, and save the heatmaps as “.png”.

NOTE: At this stage, the binding free energy (ΔG) for the lowest-energy pose (Mode 1) of each protein–ligand–pocket complex is extracted directly from the AutoDock Vina log files. These ΔG values serve as the basis for downstream thermodynamic analysis, where ΔG will be converted into dissociation constants (Kd) using the standard relationship:

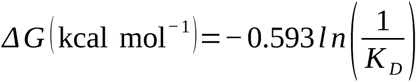

5.7. Verify that all heatmaps display proteins, drugs, and affinity values correctly, check clustering and orientation, and ensure titles and legends are accurate.

### 6. 3D Affinity Landscape Visualization and Kd conversion

NOTE: This workflow assumes that all ADME descriptors are generated externally and imported into R as a structured table. After descriptor integration and heatmap visualization, the script also produces interactive 3D affinity landscapes and converts docking-derived binding energies (ΔG) into temperature-dependent dissociation constants (Kd).

6.1. (Automated) Load required additional packages (e.g., **plotly**, **viridis**, **htmlwidgets**). Create **df_plot** from **df_results** by parsing pocket numbers with **parse_number()** and converting protein to an ordered factor via **mutate()**. Filter physiologically relevant affinities using **filter(affinity > −20 & affinity < 0)** to create **df_filtered2**.

6.2. (Automated) Construct the 3D scatter and surface. Define axis domains and a flat reference plane; then build an interactive 3D scatter with **plot_ly()** and **add_markers()**, by adding a semi-transparent surface via **add_surface()**. Customize 3D plot axes/legend and the color palette to map compounds to colors and text for hover details.

6.3. (Automated) Identify the top-20 drugs by mean affinity using **aggregate().** Subset the data to create **df_top**, and add traces per drug with **add_trace()** (larger markers for emphasis). Render the interactive figure and export it as “.html” with **saveWidget()**.

NOTE: At this stage, compounds are ranked by their overall mean binding affinity and presented in a dedicated Plot. This provides a clearer comparative landscape without explicitly subdividing results by protein or pocket.

6.4. (Automated) Export filtered docking results (**df_filtered**) by writing the affinity table to “dock_results.csv”, ensuring that downstream thermodynamic calculations operate on a clean input file. Compute temperature-dependent dissociation constants (Kd) by reading the “.csv”, expanding the data across a physiological temperature range (37 to 42 °C). Create and save the resulting long-format dataset (**long_df**) to “kd_transition.csv” for subsequent plotting after applying the thermodynamic relationship inside **mutate()**.

NOTE: The temperature-dependent Kd dataset (37-42°C range) is saved to “kd_transition.csv” in each protein subdirectory as an optional dataset for curation (Section 6.5). This temperature range is not used in the main script analyses. All efficiency metrics (pKd, LLE, FQ) are calculated using the fixed temperature of approximately 310.15 K (37°C, physiological temperature), where Kd represents the following relationship:

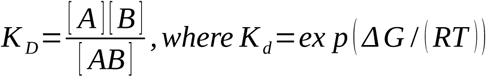

6.5. (Automated) Generate an optional quality control line plot of Kd vs temperature with **ggplot()** and **geom_line()** to inspect temperature dependence.

### 7. ADME Integration and Ligand Efficiency Scoring

7.1. (Automated) Define **parse_vina_log()** to read each “.log”, extract ranked modes with their affinities and RMSD columns, convert to a tibble, and aggregate all logs to create **df_modes**. Summarize per protein–drug–pocket with **group_by()**, **arrange()**, and **summarise()** to obtain **top1**, **top2**, and RMSD values (**rmsd1_lb**, **rmsd1_ub**) for pose-quality control as **df_poseqc**.

7.2. (Automated) Read the “adme_features_for_efficiency.csv”, normalize column names with **janitor::clean_names()**, and generate a standardized merge key using a custom **norm_key()** function (lowercase, ASCII transliteration, strip punctuation) to create **adme2** with **drug_key** column.

NOTE: Object **adme2** is necessary as an intermediate dataset that will be used for: aggregation to create **adme2_u** (merging with docking data in step 7.3), and drug-likeness rule calculations and BOILED-Egg plots (section 8).

7.3. (Automated) Aggregate ADME features per compound and collapse duplicate ADME entries. Produce **adme2_u** to merge docking and ADME data.

7.3.1. (Automated) Compute efficiency metrics, and create **df_min2** with **drug_key** from **df_min** and left-join docking summaries to ADME using **left_join()**.

7.3.2. (Automated) Compute thermodynamic and ligand-efficiency metrics (Kd, pKd, LE, LLE, FQ) in **mutate()** to create **df_eff** considering the relationships in the Supplementary material (File 1, section 1.1).

7.4. (Automated) Use **df_eff**, which contains per-compound: **min_affinity**, **HAC**, **logP_sel**, **pKd**, **LE**, **LLE**, and **FQ**. Filter compounds by LLE range (LLE *≥−* 7 *∧* LLE *≤* 7) to create **df_plot**. Plot LLE vs pKd using **ggplot()** with reference lines, and color by protein, then save the figure as “.png”.

7.5. (Automated) Render LE distributions per protein with **geom_violin()** and **geom_jitter()**. Save the complete dataset (**df_eff**) with all compounds and descriptors as “All_docked_compounds_{PROTEIN_NAME}.csv”.

NOTE: This file contains all compounds (not filtered) and is used for downstream multivariate analysis. The CSV includes columns: **tpsa**, **hbd**, **hba**, **rot_bonds**, **n_hetero** (these will be calculated from SMILES via RCDK in section 8).

7.6. (Automated) Create **df_filtered3** by ranking compounds from **df_eff** by LLE (**arrange(desc(LLE))**) and selecting the top-20 (**slice_head()**). Save it as “Best_docked_compounds_{PROTEIN_NAME}.csv”. Overlay points meeting “pKd ≥ 7 & LLE ≥ 4” using a different shape/color (shape = 21, fill = “red”) to clearly mark ideal hits in the LLE vs pKd plot.

NOTE: The **df_filtered3** contains only the top-ranked compounds by ligand efficiency, including the pocket information corresponding to the best binding affinity for each compound. The corresponding CSV includes columns: **drug**, **protein**, **pocket**, **min_affinity**, **pKd**, **hac**, **logP_sel**, **LLE**, and **FQ**. If LLE calculation fails, ensure HAC > 0 and logP values are not “NA”.

### 8. Detailed Pharmacokinetics Descriptors Calculations

NOTE: Within the script, the preferred logP value (**logP_sel**) is computed by automatically selecting the first available entry among several possible logP columns (e.g., XlogP3, WLOGP, MLOGP, iLOGP, or SwissADME consensus values). Drug-likeness rules (Veber^65^, Ghose^69^, Egan^70^, and Lipinski^71^) and BOILED-Egg absorption/BBB classifications^72^ apply directly to the descriptors already present in the **adme2** dataframe, derived from “adme_features_for_efficiency.csv”. Established drug-likeness rules are detailed in the Supplementary Material (File 1, section 1.1)

8.1. (Automated) Load required libraries (e.g., **forcats** and **rcdk**). Identify available LogP columns and recreate a single standardized field **logP_sel** by prioritizing candidates with **intersect()** and selecting the first non-missing value for **adme2**.

NOTE: Because LogP algorithm-specific estimates differ in numerical scale, mixing them within the same dataset can affect descriptor-based analyses (e.g., PCA or clustering) and introduce artificial variance unrelated to chemical properties. The provided script selects the first available LogP entry by default, but this can be modified if uniform descriptor selection is required.

8.2. (Automated) Compute rule-specific logical flags (**ghose_pass**, **egan_pass**, **veber_pass**, **lipinski_pass**) using **mutate()** and logical tests on existing ADME fields (**mw**, **mr**, **tpsa**, **hac**, **logP_sel**, etc.). Calculate the Ghose and Egan rules immediately, since all required descriptors are available from the entry “.csv” file. For Veber and Lipinski rules (which require rotatable bonds, HBD/HBA from SMILES), initialize placeholders (NA) to be filled after step 8.3.

8.3. (Automated) If SMILES are provided, parse and sanitize them using helper functions. Validate and prepare molecules, then evaluate descriptors to obtain **rot_bonds**, **hbd**, **hba**, **fsp3**, and **n_hetero** (heteroatom count) with **rcdklibs**. Bind these to **adme2** and add **tpsa**, **hbd**, **hba**, **rot_bonds**, and **n_hetero** to **df_eff** via **left_join()** using **drug_key**.

NOTE: When SMILES strings are not available, values remain NA without affecting subsequent analyses. If RCDK fails, confirm that **rcdklibs** is installed and Java is properly configured.

8.4. (Automated) Re-evaluate **veber_pass** and **lipinski_pass** using **mutate()** with the newly attached descriptors, producing final pass/fail flags for each rule. Determine **egg_hia** (intestinal absorption) and **egg_bbb** (blood-brain barrier penetration) using TPSA and WLOGP/logP_sel.

NOTE: HIA-positive (white region) aggregates compounds with TPSA *≤* 131.6 *A* °^2^ *∧−* 0.7 *≤* logP *≤* 6. BBB-positive (yolk region) aggregates compounds with TPSA *≤* 90 *A* °^2^ *∧−* 0.7 *≤* logP *≤* 6.

8.5. (Automated) Produce summary metrics with **summarise()** (percentages passing each rule) in **adme_summary**. Create a per-compound compliance matrix (**rule_df**) and aggregate a compliance bar plot for **pass_df**.

8.6. (Automated) Generate BOILED-Egg plots (TPSA vs WLOGP or logP_sel), including HIA/BBB thematic layers using **annotate()** and **geom_point()**. Create a correlation heatmap for numeric ADME properties (**MW**, **MR**, **TPSA**, **logP_sel**, **HAC**) using Spearman correlation (**corr_mat**, **corr_long**) and **geom_tile()**.

NOTE: The BOILED-Egg plots, compliance matrix, compliance bars, and correlation heatmap generated in this section are displayed only in the R environment and are not saved as files. These visualizations use global ADME data (identical across all proteins) and are regenerated for each protein during script execution. They can be viewed in the RStudio Plots panel during or after script completion. If HIA/BBB predictions (BOILED-Egg), drug-likeness compliance, or ADME correlations are not properly calculated or visualized, verify that: (1) **adme2** contains all required columns (Section 8.2); (2) SMILES strings are available; (3) there are no missing values (NA) in critical columns.

8.9. (Automated) If docking results (**df_eff**) exist, merge ADME descriptors by **drug_key** (via **left_join()**) and produce integrated analyses such as LLE vs logP_sel and pKd vs TPSA using **ggplot()** to inform compound prioritization.

NOTE: ADME correlation analysis and drug-likeness rule evaluation (Ghose, Egan, Veber, Lipinski, BOILED-Egg) do not automatically filter the output “.csv” files. Instead, visualization objects (correlation matrix, per-compound compliance matrix, compliance bar plot, BOILED-Egg plot) are displayed in the R environment for user inspection. Correlation analysis identifies multicollinearity among ADME descriptors, informing downstream multivariate analyses (e.g., PCA, clustering) by guiding feature selection. In this case, highly correlated descriptors may be combined or weighted in PCA to reduce dimensionality while preserving variance, while drug-likeness compliance enables manual compound prioritization. Users should review these visualizations to manually prioritize candidates before proceeding to multivariate analysis.

### 9. Multivariate Analysis and Visualization

NOTE: At this stage, use the automated R script “Multivariate_analysis.R” for comprehensive multivariate analysis and visualization of docking results across multiple proteins. This R script is provided in the Supplementary Material (script 3). The script reads “Best_docked_compounds_*.csv” and “All_docked_compounds_*.csv” files generated by “Main_analysis.R” (see Sections 7.5 and 7.6) and produces dimensionality reduction plots, clustering analyses, and interactive 3D visualizations.

9.1. Check if “Best_docked_compounds_*.csv” and “All_docked_compounds_*.csv” files are located in each protein directory (e.g., **∼/Vina_Workspace/receptors/{PROTEIN_NAME}/**) before proceeding.

NOTE: If “All_docked_compounds_*.csv” files are missing or lack **HAC/MW** descriptors, the 3D molecular descriptor plots will be skipped. Optional descriptors are used if available for calculating the Structural Complexity Index (SCI) in section 9.10.

CRITICAL STEP. Open Rstudio. Go to File -> Open File. Find “Multivariate_analysis.R” inside **/Desktop/Vina-SBVS-LBVS_ADME/Vina_Scripts** and load it. Install required R packages if not already installed, by running the following command in the R console:

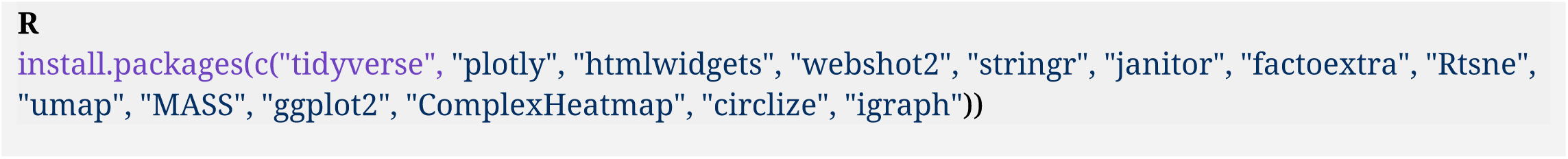

9.2. Execute the entire script in the R console. Select all text inside “Multivariate_analysis.R” script with Ctrl+A and press Ctrl+Enter (or click “Run”).

9.3. (Automated) Load required R packages. Set working directory based on **VINA_WORKSPACE** variable. Automatically detect all “Best_docked_compounds_*.csv” and “All_docked_compounds_*.csv” files in protein subdirectories using **list.dirs()** and **list.files()**.

9.4. (Automated) Read and combine all CSV files using **read_with_protein()** and **read_all_compounds()** functions. Extract protein names from filenames and automatically extract virus/identifier (first part before separator). Map **min_affinity** to **affinity** (standardized name), and handle pocket information if available.

NOTE: Protein names can use any separator (_, -, space, .), and the script will extract the identifier automatically for grouping. If no clear identifier is found, virus grouping is set to NA, and the script proceeds with protein-only grouping. If any CSV file lacks the required **affinity** or **drug** columns, that file is skipped with a warning. The pocket field is used only for visualization in interactive 3D plots in hover text. If the CSV does not contain pocket, the plots work normally.

9.5. (Automated) Filter data to retain physiologically relevant affinities (affinity ≤ −9 kcal/mol) used to create **df_filtered**. Generate interactive 3D docking landscapes using **plot_ly()**: global 3D plot (protein × compound × affinity); per-virus 3D plots and per-virus-per-protein 3D plots. Export all plots as “.html” (interactive) and “.png” files.

NOTE: All three 3D plots share the same axis structure: X-axis = protein, Y-axis = compound (drug), Z-axis = affinity (kcal/mol). Protein and virus information are represented through color coding (not axes). The global plot displays all protein-compound interactions from the entire dataset in a single view, where each point represents a specific protein-compound interaction. This plot is colored by virus (when available) or by protein (if virus grouping is unavailable). Per-virus plots display only interactions involving proteins from a specific viral species, colored by protein to enable cross-protein comparisons within that virus. Per-virus-per-protein plots display only interactions for a specific virus-protein combination (single protein position on X-axis), colored by compound. Pocket information appears as marker symbols when available. When virus grouping is unavailable, per-virus plots are not generated.

9.6. (Automated) Prepare affinity matrix (proteins × compounds) by aggregating the minimum affinity per protein–ligand pair. The matrix is constructed as: **mat[protein, compound] = min(affinity)** for each protein–compound pair, with missing values filled as 0. Convert to a numeric matrix using **as.matrix()**.

9.7. (Automated) Perform Principal Component Analysis (PCA) using **prcomp(mat, scale. = TRUE)**. Generate PCA biplot and individual plots colored by virus and/or protein using **factoextra** package. Save plots as “.png” files.

NOTE: PCA reduces dimensionality by identifying orthogonal directions of maximum variance in the affinity data, allowing visualization of protein–compound relationships in lower-dimensional space. If virus = NA for all entries, virus-colored PCA plots are skipped, and only protein-colored plots are generated.

9.8. (Automated) Perform dimensionality reduction analyses: UMAP (Uniform Manifold Approximation and Projection) using **umap()** with **n_neighbors = 10**, **min_dist = 0.2**. Plot UMAP1 vs UMAP2 colored by virus and shaped by protein; t-SNE (t-distributed Stochastic Neighbor Embedding) using **Rtsne()** with **perplexity = 5**, **theta = 0.5**. Plot tSNE1 vs tSNE2 colored by virus and shaped by protein. Save all plots as “.png” files.

NOTE: UMAP is a non-linear dimensionality reduction that preserves local and global structure, and t-SNE is a non-linear technique that emphasizes clustering. These methods require minimum data points relative to their parameters (UMAP: **n_neighbors = 10;** t-SNE: **perplexity = 5**). If insufficient data is present or an error occurs, these analyses are skipped with a warning message, and the script continues execution.

9.9. (Automated) Extract molecular descriptors from the “.csv” files and calculate the Structural Complexity Index (SCI). Prepare compound × protein affinity matrix (**mat_compounds**) by aggregating minimum affinity per compound–protein pair, then pivot to matrix format. Create **meta_compounds** with descriptors and SCI and ensure alignment of compounds between matrix and metadata using **intersect()**.

NOTE: If “All_docked_compounds_*.csv” files are missing or lack **HAC/MW** columns, Section 9.13 (3D plots colored by molecular descriptors) is skipped with a warning message. SCI is calculated using an approximate custom relationship presented in the Supplementary Material (File 1, section 1.1).

9.10. (Automated) Generate a PCA 3D plot colored by Structural Complexity Index. Perform PCA on **mat_compounds**, extract PC1, PC2, PC3, and color points by SCI values (red = simple scaffolds, blue = complex polycyclic/heteroatom-rich structures). Export as “.html” and “.png”.

NOTE: If SCI cannot be calculated due to missing optional descriptors, the code falls back to **HAC** for coloring. **HAC** reflects only molecular size, whereas SCI combines size, heteroatom richness, and rigidity, so the fallback may provide less discrimination between simple and complex structures.

9.11. (Automated) Generate t-SNE 3D plot colored by molecular weight. Perform 3D t-SNE on **mat_compounds**, extract tSNE1, tSNE2, tSNE3, and color points by **MW** values. Export as “.html” and “.png”.

9.12. (Automated) Perform Linear Discriminant Analysis (LDA) if multiple virus groups are available **(length(unique(virus)) > 1)**. Apply **lda()** for classification, extract LD1 and LD2, and plot colored by virus and shaped by protein. Save as “.png”.

NOTE: LDA is a supervised dimensionality reduction technique that maximizes separation between predefined groups (e.g., viruses). LDA requires at least 2 distinct virus groups. The script checks if more than one unique virus value exists (**length(unique(meta$virus)) > 1**); if only one virus group exists, the LDA plot is skipped. The current implementation does not explicitly filter NA values, so if one virus group plus NA values are present, LDA may be attempted, but could fail if insufficient non-NA groups exist.

9.13. (Automated) Generate distribution plots: a violin plot as an affinity distribution per virus using **geom_violin()** and **geom_boxplot()**; a Multi-target barplot with compounds targeting multiple proteins (n_targets > 1) ranked by number of targets and mean affinity. Save all plots as “.png” files.

NOTE: The violin plot is skipped if virus grouping is not available, but the Multi-target barplot is always generated if multi-target compounds exist.

9.14. (Automated) Generate a hierarchical clustering heatmap using **ComplexHeatmap**. Prepare an affinity matrix for compounds with affinity ≤ −9 kcal/mol and aggregate values by virus-protein (or protein) and drug. Pivot to matrix format, apply **colorRamp2()** for the color scale (red-yellow-white for negative affinities), and add row annotations for virus grouping if available. Save as “.png”.

9.15. (Automated) Generate a 2D intensity map using **ggplot2**. Plot protein (x-axis) vs drug (y-axis) with point size proportional to absolute affinity and color by virus (if available). Save as “.png”.

9.16. Verify that all output files are generated in **∼/Vina_Workspace/multivariate_analysis_output/plots_3D/** subdirectory: “html” and “png” files for 3D interactive plots. In **∼/Vina_Workspace/multivariate_analysis_output/**, check: “.png” files for PCA, UMAP, t-SNE, LDA, violin, barplot, heatmap, and intensity map.

NOTE: If **webshot2** fails to convert “.html” plots to “.png” (e.g., due to missing Chrome/Chromium), the script continues execution and generates all subsequent plots. The “.html” files are saved before conversion and can be opened in a web browser. Ensure Chrome/Chromium is configured for headless rendering or run **webshot2::install_phantomjs()** if using PhantomJS.

## REPRESENTATIVE RESULTS

Following the completion of all workflow steps, multivariate analysis outputs are generated and stored in ∼/Vina_Workspace/multivariate_analysis_output/. These multivariate results integrate and compare data from Sections 4 to 8, including: (i) docking-derived binding affinities (ΔG) and pocket-specific interactions from AutoDock Vina log files (Section 4); (ii) binding affinity heatmaps and minimum ΔG values per protein–ligand pair (section 5); (iii) 3D affinity landscapes, dissociation constants (Kd), and temperature-dependent thermodynamic analyses (Section 6); (iv) ADME-integrated ligand efficiency metrics (pKd, LLE, LE, FQ) and efficiency-based compound rankings (Section 7); and (v) pharmacokinetic descriptors, drug-likeness rule compliance (Lipinski, Veber, Ghose, Egan), and BOILED-Egg absorption/BBB predictions (section 8). This integrated comparison enables comprehensive chemical and biological inference and interpretation by revealing relationships between binding thermodynamics, ligand efficiency, and pharmacokinetic properties across multiple viral protein targets used in this study. The complete workflow is summarized in the following flowchart (Figure 1).

### Binding Affinities and Main Inhibitors by Viral Protein

Docking experiments identified, for each viral protein, candidate compounds exhibiting notably high binding affinities (very negative ΔG values, indicative of favorable interactions). Table 1 summarizes the top 10 performing compounds for each viral protein, with average binding affinities (ΔG) calculated across all binding pockets and the five Flavivirus species. Reported affinity values correspond to the most favorable predicted ΔG used for pKd estimation. The completely refined data set is in the Supplementary Material (File 3). The compounds’ ligand-lipophilicity efficiency (LLE) profiles will be further discussed, providing a key criterion for prioritization. ADME features used to calculate LLE can be assessed in the Supplementary Material (File 4). [Place **Table 1** here]

**Table 1.** Top-performing compounds per viral protein target. Displayed ΔG values (kcal·mol⁻¹) are mean protein-related affinities computed across all Flavivirus species and are shown only for compounds ranking among the top 10 for each protein type. The “Main contributing species” column identifies the viruses that most influenced these averages; individual per-virus contributions may not be reflected if the corresponding values did not rank among the top 10 compounds for the analyzed protein.

In addition to the averaged binding affinities, this workflow also examined the most favorable individual interactions observed for each viral protein. Table 2 lists the top 15 compounds based on their minimum predicted binding free energy (ΔGₘᵢₙ) values among all binding pockets for the respective protein targets. These values highlight compounds capable of achieving strong interactions within at least one binding site, thus complementing the averaged affinity profiles. [Place Table 2 here]

**Table 2.** Top-performing compounds per viral protein target, ranked by minimum binding affinity (ΔGₘᵢₙ, kcal·mol⁻¹) observed among all pockets. Displayed values correspond to the lowest predicted ΔG identified for each top 15 compounds across all Flavivirus species.

Several results are particularly noteworthy. The NS3 protease was identified as a target for one of the most potent ligands: temoporfin, a porphyrinic compound, which displayed a binding energy of approximately –11 kcal/mol, corresponding to a Kd near 1 nM (pKd ≈ 9). This extraordinary affinity suggests that temoporfin binds very stably within the active site of NS3, potentially “docking” between the loops that interact with the NS2B cofactor and thereby blocking the proteolytic activity essential for viral polyprotein maturation^73^. However, temoporfin has a logP of ∼ 8.8, reflecting its extremely lipophilic nature^74^. Its lipophilic efficiency (LLE) is nearly zero, indicating that much of its high binding affinity may result from non-specific hydrophobic interactions—a cautionary signal regarding its pharmacological profile. In other words, although temoporfin is potent *in vitro*, its solubility is likely limited, and it may preferentially partition into cell membranes, necessitating special formulations (e.g., by using lipid vehicles for administration). Therefore, temoporfin exemplifies the classic trade-off between potency and pharmacokinetic properties: it leads in NS3 binding affinity but may require optimization—or replacement with less lipophilic analogs—to be viable as a systemic antiviral.

The viral polymerase NS5 highlighted a different type of inhibitor: sofosbuvir, a nucleoside analog approved for hepatitis C virus (HCV), exhibited strong predicted binding in the catalytic site of NS5 with an estimated affinity ∼ −9 kcal/mol, pKd ∼ 6–7 (data not shown). Sofosbuvir is metabolized intracellularly into its triphosphate form, which competes with natural nucleotides at the polymerase; our results suggest that, even in its prodrug state, it can occupy the nucleotide binding site of NS5, consistent with its anticipated mechanism of action^75^. Unlike temoporfin, sofosbuvir is polar, containing masked phosphate groups, and demonstrates high LLE—achieving moderate potency without relying on high lipophilicity. This characteristic reinforces sofosbuvir’s potential as a repurposing candidate for flaviviruses. Indeed, previous studies have shown that sofosbuvir inhibits ZIKV replication in cell culture and murine models, reducing viral loads across multiple tissues and improving survival of infected animals^76, 77^. Accordingly, our docking data support sofosbuvir’s capacity to act as a pan-flaviviral NS5 inhibitor, given the high sequence homology among polymerases of these viruses (e.g., ∼ 80% amino acid identity between ZIKV and HCV polymerase domains)^78^.

Regarding the viral structural proteins, we observed promising binding, although generally with less extreme affinities than for the enzymatic targets. For the envelope (E) protein, several natural flavonoids emerged as good ligands (data not shown). Polyphenolic compounds such as myricetin and quercetin exhibited affinities in the range of 2–10 µM (pKd ∼ 5–5.7, ΔG reaching values below −7 kcal/mol), not only for E but also, intriguingly, for NS5. These flavonoids appear to interact at an allosteric site within the E domain, potentially stabilizing the protein in a conformation that is incompetent for viral fusion or assembly79. Simultaneously, their planar aromatic cores and hydroxyl groups, capable of forming hydrogen bonds, allow them to fit into the nucleotide binding site of NS5, suggesting a possible dual mechanism of action. The comparative analyses indicate that several flavonoids exhibited this same dual binding in more than 4 viruses, which supports other similar studies and highlights a remarkable multi-target and pan-viral potential for this class of compounds80, 81. Their favorable LLE profiles further reinforce the attractiveness of flavonoids: myricetin, for example, combines a pKd ∼ 5.7 with a modest logP (∼ 1–2), resulting in an LLE of ∼ 4, considered optimal. In other words, these compounds achieve moderate potency primarily through polar and specific interactions rather than non-specific hydrophobic contacts, which generally confer lower toxicity and good solubility. Together with evidence of multi-target activity, these properties make flavonoids such as myricetin strong lead candidates for future optimization82. Example docked poses of a subset of screened compounds in the NS5 polymerase binding site are shown in Figure 2, illustrating typical ligand orientations and predicted interactions within the conserved catalytic pockets among the viral targets.

**Figure 2.**
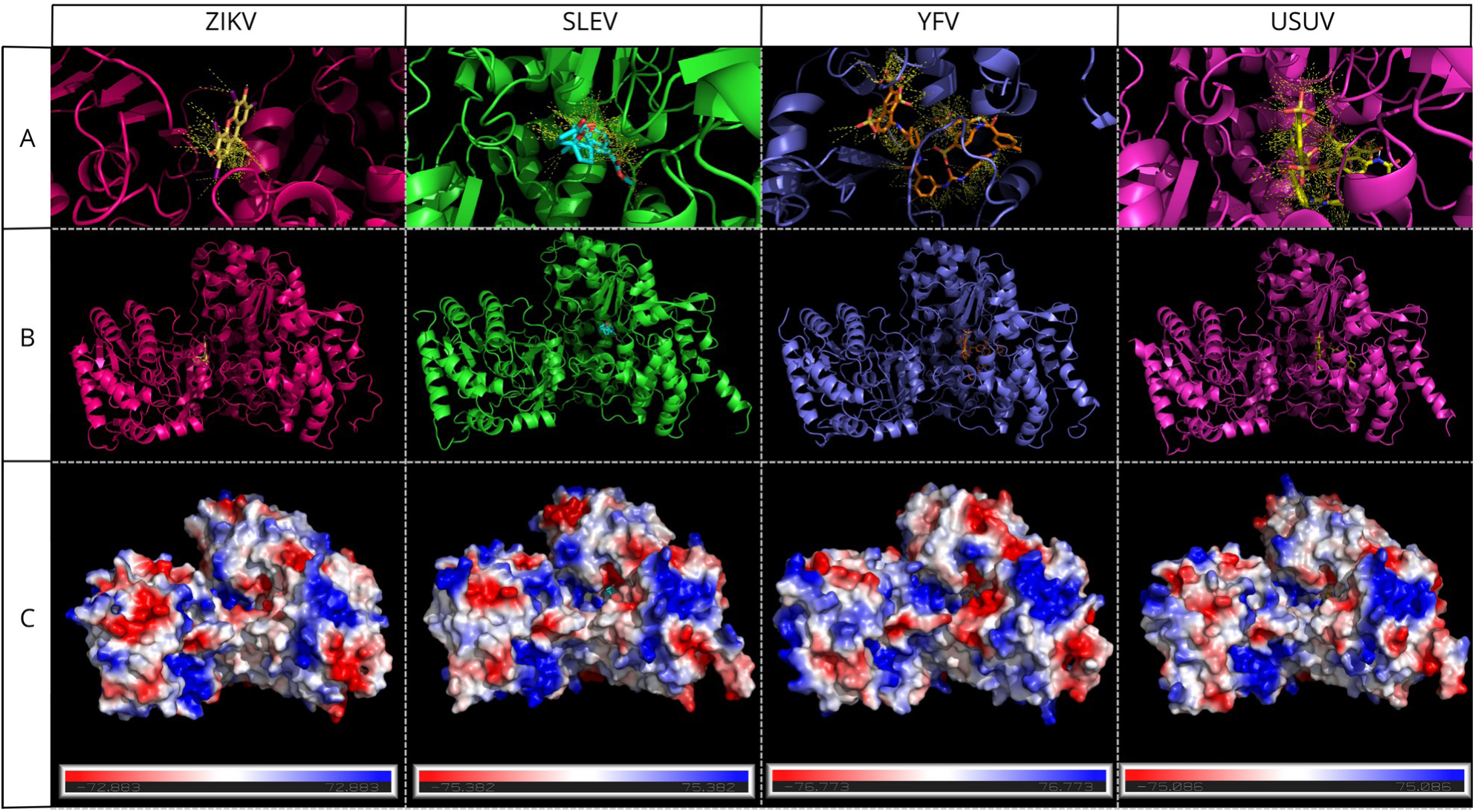
Docked poses of a subset of ligand compounds in the NS5 polymerase binding site across all Flavivirus species. (A) Cartoon representation of NS5 with selected ligands and their predicted contacts within the conserved binding pocket, shown using a 4 Å distance threshold. (B) Broader view of the corresponding NS5–ligand complex. (C) Electrostatic surface of the corresponding NS5 generated using APBS (Adaptive Poisson-Boltzmann Solver), highlighting the distribution of charges within the binding site.

For the capsid (C) protein, our docking results suggest that both the RNA binding site and capsid dimers can accommodate polyanionic compounds. Aurintricarboxylic acid (ATA), a well-characterized nucleic acid inhibitor, exhibited relatively high affinity (approximately low µM) for the capsid (data not shown). ATA is a large, highly charged molecule (tricarboxylated) known to interfere with protein–nucleic acid interactions. Its pKd of ∼ 5.7, combined with a very low logP (extremely polar), results in a high LLE, indicating that it achieves reasonable potency without relying on hydrophobic interactions. This compound has demonstrated broad-spectrum antiviral activity *in vitro* against human immunodeficiency virus (HIV) and other viruses, acting primarily on nucleoprotein processes ^83, 84^. However, despite its promising predicted affinity, ATA did not pass the ADMET filters applied. This is consistent with the fact that it has not been approved by the FDA for antiviral use, and, to our knowledge, no New Drug Application (NDA) has been granted for this indication. Consequently, it was not included in the refined docking output table (File 3, Supplementary Material) and derivative plots. Nonetheless, the identification of ATA as a capsid ligand supports the feasibility of targeting capsid functions—such as RNA encapsidation or nucleocapsid assembly—and highlights the potential value of developing ATA analogs with improved selectivity and bioavailability, given that highly anionic molecules generally face challenges crossing cell membranes.

The small membrane proteins (prM/M) and some nonstructural proteins (NS2A, NS2B, NS4A, NS4B) generally exhibited docking affinities dominated by highly hydrophobic compounds, consistent with the non-polar character of their environments. For example, niclosamide—an established antiparasitic drug that acts by uncoupling proton gradients—emerged among the top candidates for M and NS4B. Niclosamide is strongly hydrophobic (logP ∼ 5) and effectively occupies predicted lipid-accessible pockets in these membrane proteins, with ΔG values in the range of −7 to −8 kcal/mol. Its LLE of approximately 3–4 indicates moderate potency (µM) despite relatively high lipophilicity^85^. Interestingly, niclosamide-related compounds have also been reported to inhibit the flavivirus NS2B–NS3 complex^86^, possibly via allosteric destabilization of the cofactor–protease interaction. Furthermore, independent studies have demonstrated broad antiviral activity for niclosamide, including against flaviviruses, partially attributed to disruption of acidic organelles (endosomes) and other pleiotropic mechanisms^87^.

In the present context, its consistent performance across multiple membrane targets suggests potential interference with key viral processes, such as fusion or replicative complex assembly, that rely on these proteins.

Overall, each viral protein yielded at least one candidate compound with affinity in the micromolar range or better, supporting the effectiveness of the multi-target virtual screening approach. However, these results also underscore that binding affinity alone is insufficient for prioritization. In this context, LLE analysis proved invaluable for filtering out or deprioritizing “false positives” -- e.g., very non-polar compounds with apparently optimal binding energies, but prone to pharmacokinetic problems. In contrast, compounds with moderate affinity but more balanced chemical profiles emerge as more promising candidates for further development.

### Compound Performance by Virus and Intervirus Comparison

In addition to analyzing individual targets, we evaluated the overall performance of compounds across all proteins within each virus. For each of the five Flaviviruses, we identified a subset of compounds that consistently exhibited strong binding across multiple viral proteins. Table 3 summarizes, for each virus, ten compounds with the best averaged performance — i.e., those achieving the highest number of favorable interactions across the virus’s various targets. The reported affinity values represent approximate averages or representative ΔG ranges reflecting the compound’s binding across different targets, accompanied by the main proteins responsible for these high-affinity interactions. [Place **Table 3** here]

**Table 3.** Top-performing compounds for each Flavivirus species. Displayed ΔG values (kcal·mol⁻¹) are shown as averages only for compounds ranking among the top 10 for each virus. The “main targets” column lists proteins with high contribution to the average affinities across all viruses, even if corresponding values are not displayed.

To complement the average-binding analysis, the most favorable individual interactions of compounds across all protein targets for each virus were also computed in this workflow. Table 4 lists the top 15 compounds per Flavivirus species based on their minimum predicted binding free energy (ΔGₘᵢₙ) observed among all proteins. This approach provides an alternative perspective on overall antiviral potential. [Place Table 4 here]

**Table 4.** Top-performing compounds for each Flavivirus species. Displayed ΔG values (kcal·mol⁻¹) correspond to the minimum predicted binding affinity (ΔGₘᵢₙ) observed across all protein targets and are shown only for compounds ranking among the top 15 for each virus.

To contextualize these rankings within the broader energetic landscape, we compared the full distribution of docking affinities for each virus in a violin plot (Figure 3). All five species display similar median values (≈ –6 kcal·mol⁻¹), indicating comparable global binding propensities and supporting the robustness of the dataset. However, ZIKV and SLEV exhibit noticeably deeper high-affinity tails (reaching ≲ –10 kcal·mol⁻¹), consistent with their prominent contribution to high-affinity clusters in the multi-target analyses. In contrast, USUV and YFV show narrower distributions, suggesting more restricted ligand recognition, while WNV presents an intermediate spread with both strong binders and a concentration of mid-range interactions. These distributional differences inform virus-specific interaction biases that help explain the compound-level trends discussed below.

**Figure 3.**
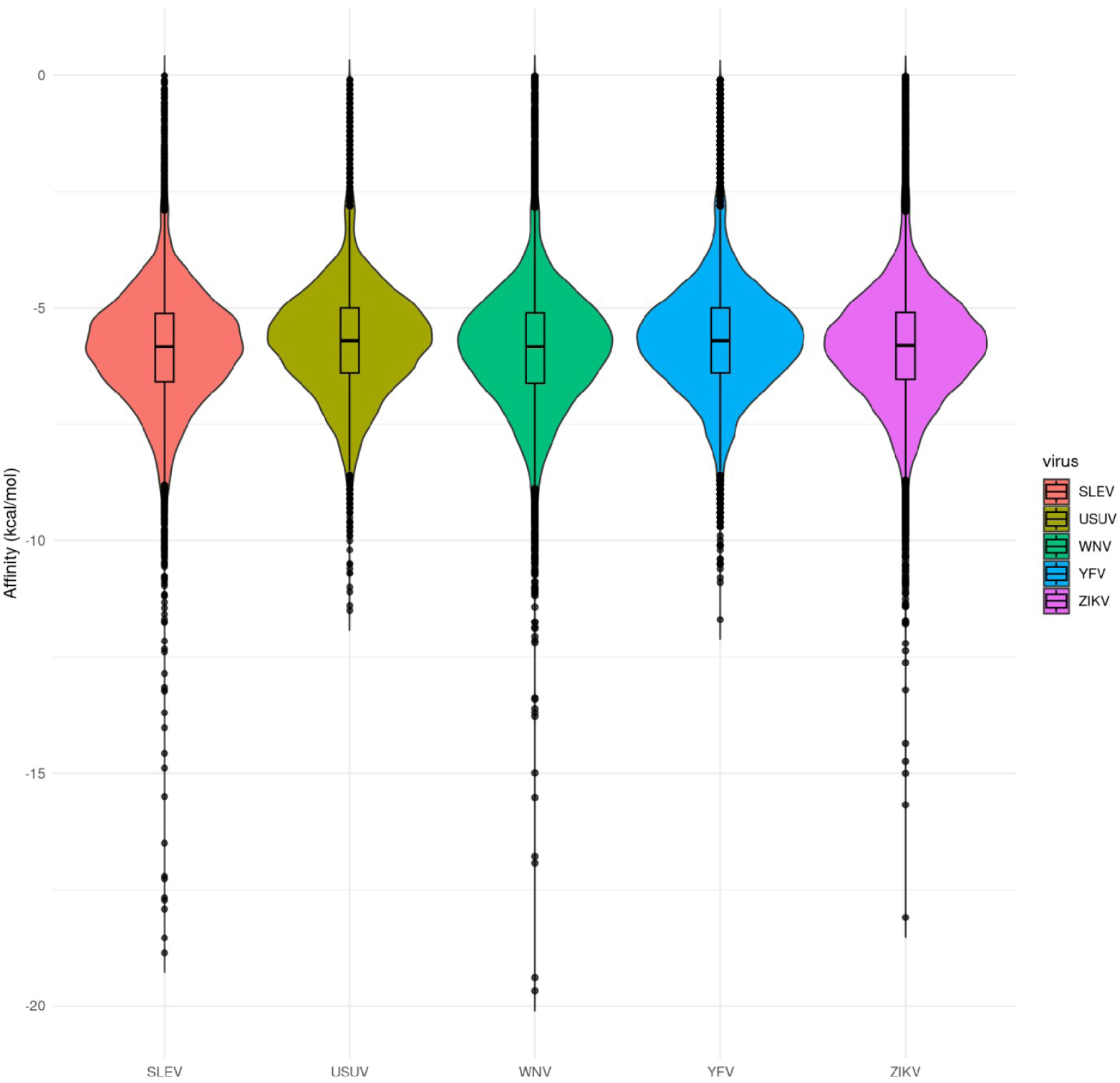
Violin plots displaying the distribution of docking affinities (kcal/mol) for all compounds across the proteomes of SLEV (red), USUV (dark green), WNV (light green), YFV (light blue), and ZIKV (pink), with boxplots showing median and interquartile range.

A prominent trend is the repeated identification of temoporfin among the top-performing ligands across Flavivirus species, alongside other compounds, such as myricetin, that exhibited high predicted affinities in our broader dataset. Temoporfin consistently ranked among the strongest inhibitors of NS3 (and the NS2B–NS3 complex) in ZIKV, USUV, YFV, SLEV, and WNV, highlighting the high conservation of the NS3 protease active site among these flaviviruses. This result aligns with prior experimental findings demonstrating its ability to inhibit ZIKV replication both *in vitro* and *in vivo*, as well as its activity against other flaviviruses^88^. In practical terms, this raises the possibility that a single drug — or a temoporfin-derived prototype — could suppress viral polyprotein cleavage across multiple pathogens, effectively blocking early stages of the replicative cycle.

Myricetin, in turn, exhibited moderate predicted binding across the analyzed viruses, generally due to its dual affinity for both the NS5 polymerase and the E glycoprotein. Similar flavonoids, such as quercetin and luteolin, showed comparable binding patterns. Notably, for YFV, myricetin approached the predicted affinity of temoporfin, suggesting that, in some viruses, an NS5/E inhibitor can be as relevant as an NS3 inhibitor when considering multi-target templates.

Aurintricarboxylic acid (ATA) was initially identified among YFV’s top three ligands, but it was excluded from further consideration due to ADMET restrictions. Its apparent prominence may reflect a particular feature of the YFV capsid enhancing ATA’s relative affinity, or stochastic variation near the selection threshold observed in other viruses. Overall, the consistent performance of myricetin and temoporfin across all five genomes highlights two preferential antiviral axes — NS5/E and NS3, respectively — supporting a broad-spectrum therapeutic strategy.

Sofosbuvir also appeared among the top compounds for SLEV and WNV, and showed consistently favorable predicted binding across other Flavivirus species. Its inclusion reinforces that nucleoside polymerase analogs, beyond being clinically validated candidates in HCV therapy, possess cross-flaviviral potential. Notably, while YFV already benefits from an effective vaccine, outbreaks continue to occur, and antiviral pharmacological alternatives could still be valuable for treating severe cases. As mentioned in the introduction, Sofosbuvir has shown significant inhibitory effects against YFV in cell culture, suggesting that combination regimens may further enhance its efficacy.

Niclosamide also exhibited moderate, but consistent predicted binding for the studied viruses, including USUV and WNV, which likely reflects the importance of membrane-associated proteins (M and NS4B) in the replication cycle of these viruses, or minor sequence variations that enhance its affinity. While not among the highest-scoring compounds, its performance remains notable given the diversity of the library screened. USUV, for instance, remains relatively underexplored; nonetheless, our data suggest that repurposing niclosamide—a low-cost, clinically safe drug — could be a promising strategy for USUV and WNV infections, given its multitarget effects, including endosomal fusion blockade and protease inhibition^86, 89^. From a comparative standpoint, no dramatic divergences were observed among the viruses regarding preferred compounds.

The modest variations observed, such as ATA appearing in YFV but not others, or niclosamide showing higher relevance in WNV, likely stem from subtle sequence differences in target proteins that influence ligand accommodation. Even so, several compounds displayed consistent multi-virus activity, suggesting that a unified therapeutic approach targeting shared structural motifs is feasible. Nonetheless, the observed variations also support the rationale for combination therapy: if a given virus shows reduced sensitivity to a single agent due to target variability, complementary compounds in a multitarget regimen could compensate for that gap.

### Ligand-Lipophilic Efficiency (LLE) and Quality of the Main Ligands

As highlighted earlier, incorporating LLE analysis was essential for distinguishing compounds that not only exhibit strong binding affinities but also possess realistic drug-like potential. Figure 4 illustrates this relationship for the Zika virus case, showing the distribution of all tested ligands according to their potency (pKd) and lipophilic ligand efficiency (LLE), with each point corresponding to a specific viral target.

**Figure 4.**
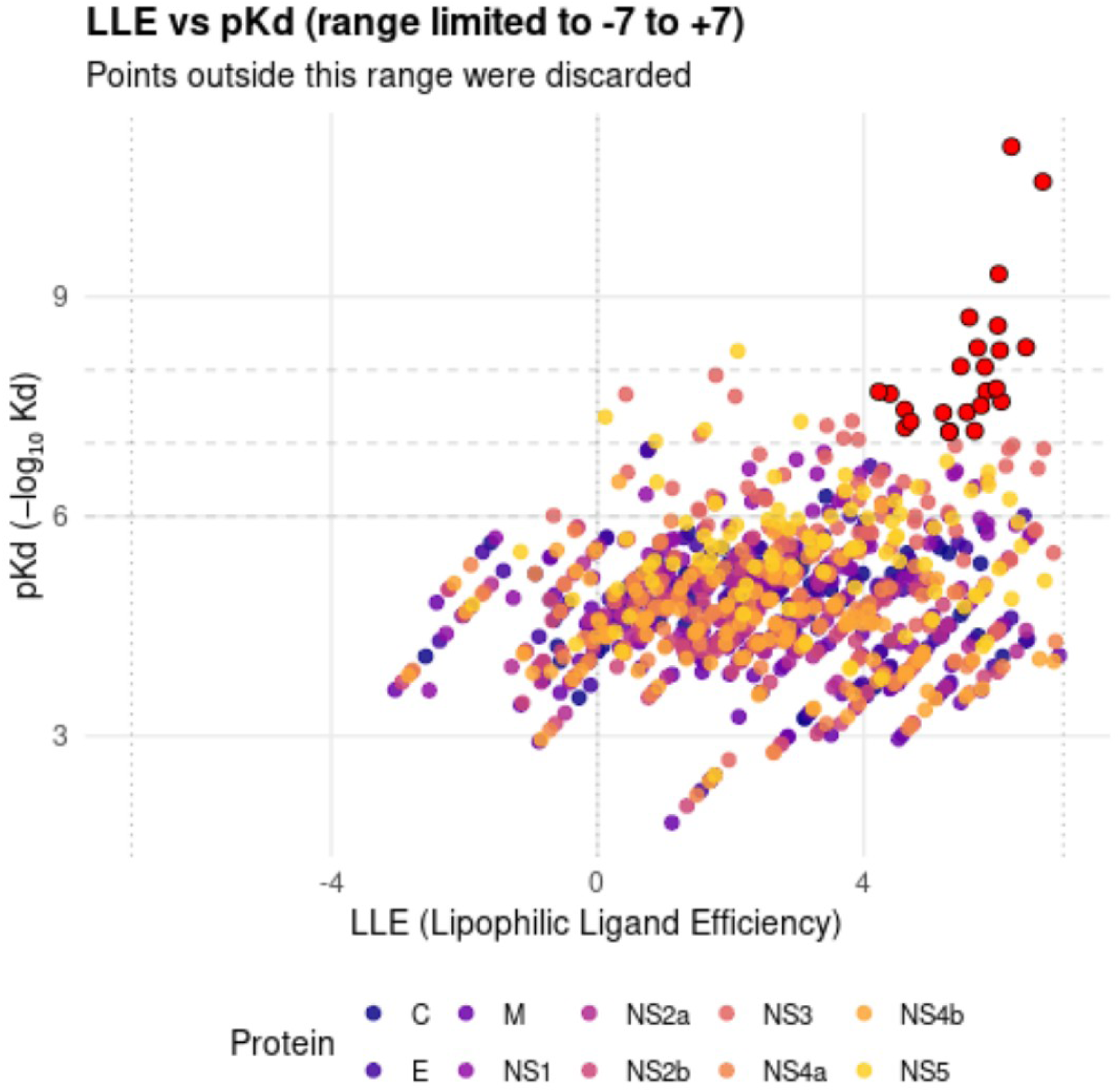
Relationship between affinity (pKd) and lipophilic efficiency (LLE) of compounds tested against Zika virus proteins. Each point represents a ligand–protein complex, colored according to the target protein. Only the LLE range between –7 and +7 is displayed, excluding outliers for clarity.

It is observed that relatively few compounds simultaneously achieved high pKd and high LLE (upper right quadrant). The most promising compounds will be those close to this region, indicating strong intrinsic potency not obtained at the cost of extreme hydrophobicity. In the case of ZIKV, a concentration of points for NS3 and NS5 is seen in the region of pKd ≥ 5 but with negative LLE, reflecting compounds like temoporfin in complex with NS3 - very potent, yet strongly lipophilic. In contrast, some points corresponding to NS5 and E (colored yellow and purple, respectively) stand out with LLE > 3: these are precisely flavonoids (like myricetin) and nucleoside analogs (sofosbuvir) that combine moderate affinity with a more polar character. Similar analysis for the other viruses revealed consistent patterns, indicating that most top hits displayed at least a reasonable LLE profile, with the notable exception of the porphyrinic compound class.

Principal component (PCA) and t-distributed Stochastic Neighbor Embedding (t-SNE) analyses (File 1, figures S1-S6 in Supplementary Material) confirmed that the compound library spans a broad and diverse region of the antiviral chemical space, supporting an unbiased and chemically heterogeneous virtual screening dataset with minimal redundancy. This chemical diversity, combined with ligand-lipophilicity efficiency (LLE) considerations, guided the prioritization of compounds such as myricetin and sofosbuvir, which balance favorable binding with suitable physicochemical properties for further development. Myricetin and similar compounds could be chemically optimized to increase potency (pKd) while maintaining or improving solubility, since they start from a favorable LLE point. This ensures that the therapeutic suggestions are aligned not only with predicted antiviral efficacy but also with drug-likeness, increasing the chances of success in experimental stages. The scatter plots containing affinity (pKd) vs lipophilic efficiency (LLE) for all flaviviruses are in Supplementary Material (File 1, S10-14).

### Cluster Analysis: Grouping Proteins and Compounds in Heatmaps

To explore global patterns in the predicted interactions, we first examined the distribution of binding affinities across all compounds and viruses. Figure 5 presents a representative heatmap summarizing these profiles, providing an at-a-glance view of major trends, with viral proteins shown as rows and a selected subset of compounds as columns. The representative subset of compounds used to generate this heatmap can be assessed in a spreadsheet “Compound_codes” inside the Supplementary Material (File 3).

**Figure 5.**
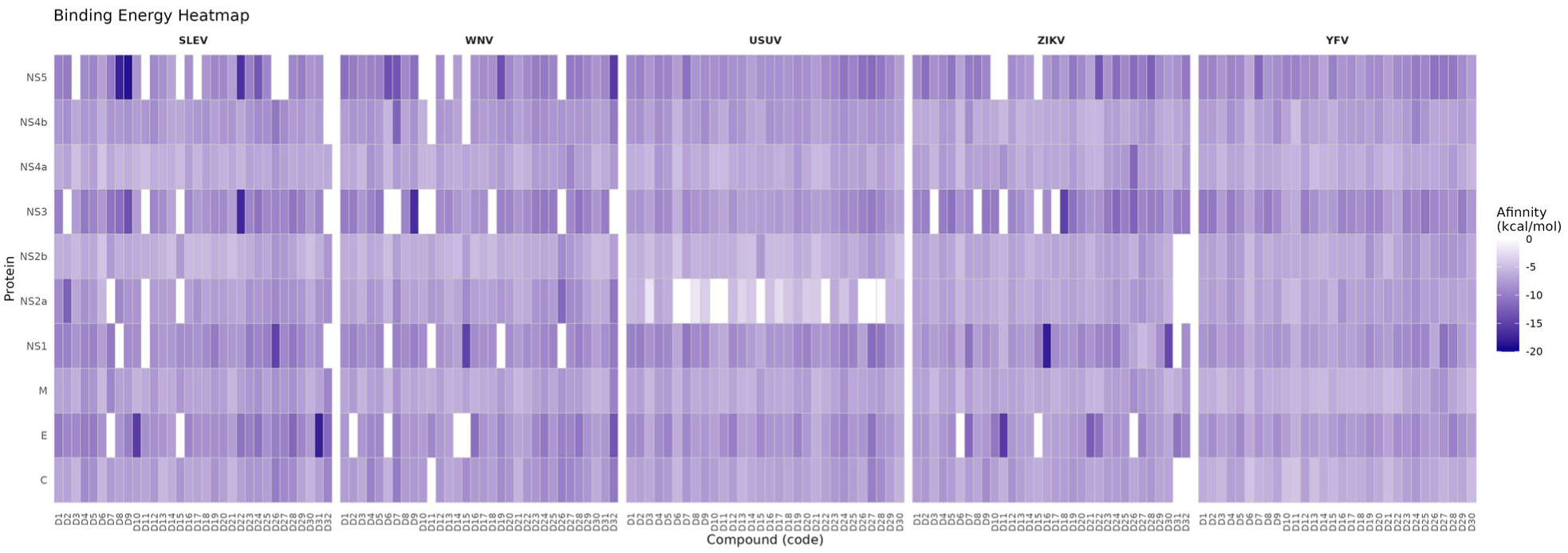
Heatmap of predicted binding affinities (ΔG, kcal/mol) for representative compounds (columns) against viral proteins (rows) for the five analyzed flaviviruses: USUV, YFV, SLEV, WNV, and ZIKV. Colors range from white (weak binding, ΔG near 0) to deep blue (strong binding, highly negative ΔG). Each code corresponding to a compound identifier can be checked in the Supplementary Material (File 3).

Another informative visualization format is presented in Figure 6, which condenses the compound–protein interaction landscape into a two-dimensional intensity map after energy-range filtering. This representation highlights the relative strength and distribution of affinities across all flaviviruses, making it easier to spot recurrent high-affinity clusters and cross-species patterns. The most prominent hotspots are associated with NS3 and NS5. These enzymes exhibit recurrent strong binding with multiple compounds, particularly Suramin, Merimepodib, and Sofosbuvir, consistent with their catalytic clefts’ broad physicochemical adaptability. Notably, Andrographolide presents a high-affinity profile for NS5, while Quercetin appears among the top hits for WNV’s NS3. Envelope (E) and NS1 proteins also display several intense interactions (e.g., Lovastatin, Ivermectin, and Digoxin). Conversely, the capsid (C) and membrane (M) proteins show fewer and weaker interactions, consistent with their compact, hydrophobic nature and reduced ligand accessibility under static docking conditions.

**Figure 6.**
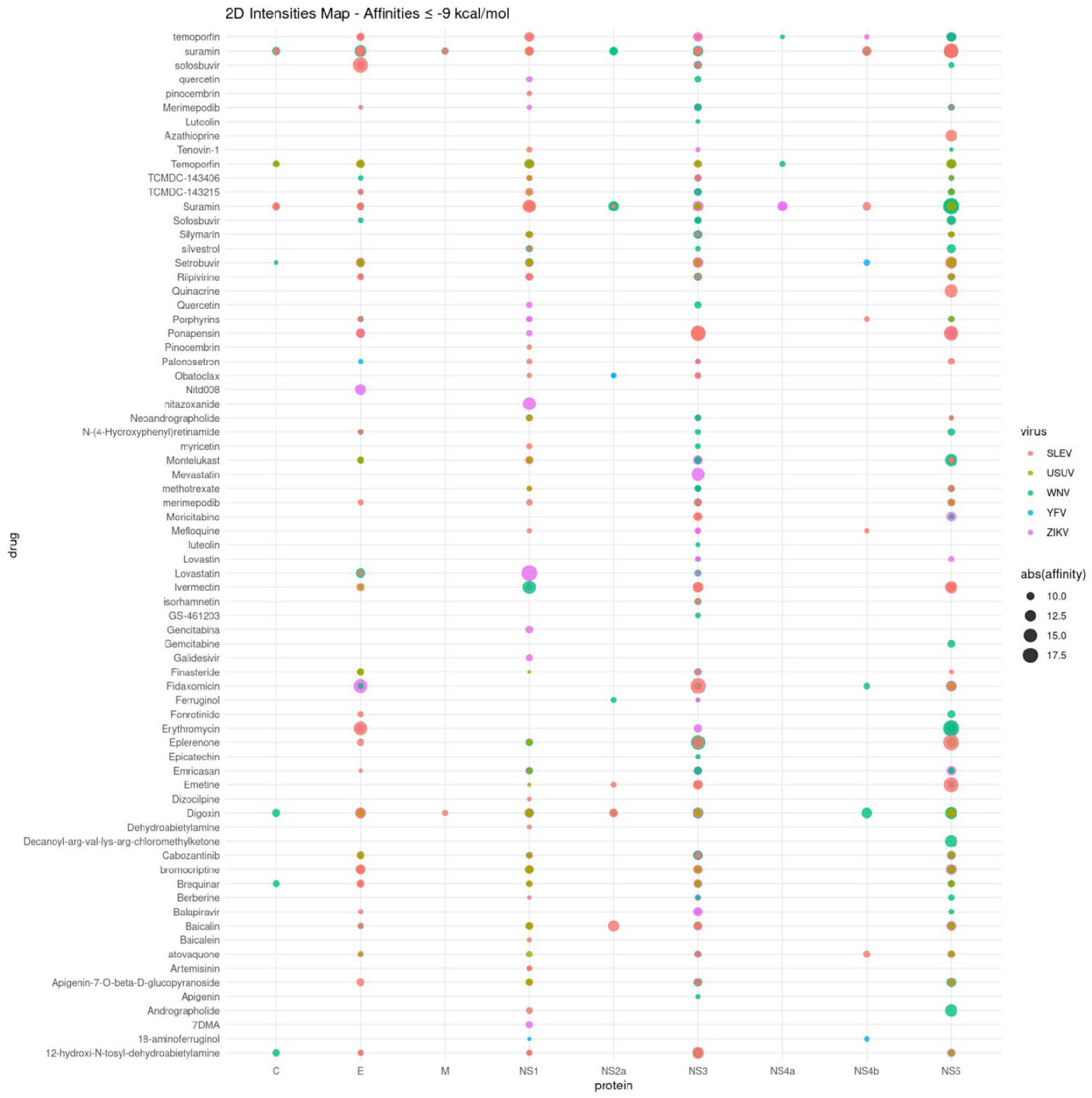
This two-dimensional intensity map summarizes the distribution and strength of binding affinities (≤ –9 kcal/mol) between all tested compounds (y-axis) and viral proteins (x-axis) across five flaviviruses (SLEV, USUV, WNV, YFV, ZIKV). Each point represents a compound–protein interaction, color-coded by virus and scaled in size according to the absolute affinity value (ΔG in kcal/mol).

The color distribution reveals a partial segregation by virus lineage, where SLEV and WNV compounds dominate NS3/NS5 hotspots, while ZIKV and YFV display additional concentration in NS4a and NS4b, respectively. USUV interactions appear more evenly distributed but of slightly lower intensity, suggesting a narrower ligand compatibility range. Building on this overview, a full hierarchical clustering analysis was conducted across the complete interaction matrix (Figure 7), providing a more detailed organization of compound–virus relationships. This approach highlights groups of compounds that share coherent biochemical features and multi-target interaction profiles.

**Figure 7.**
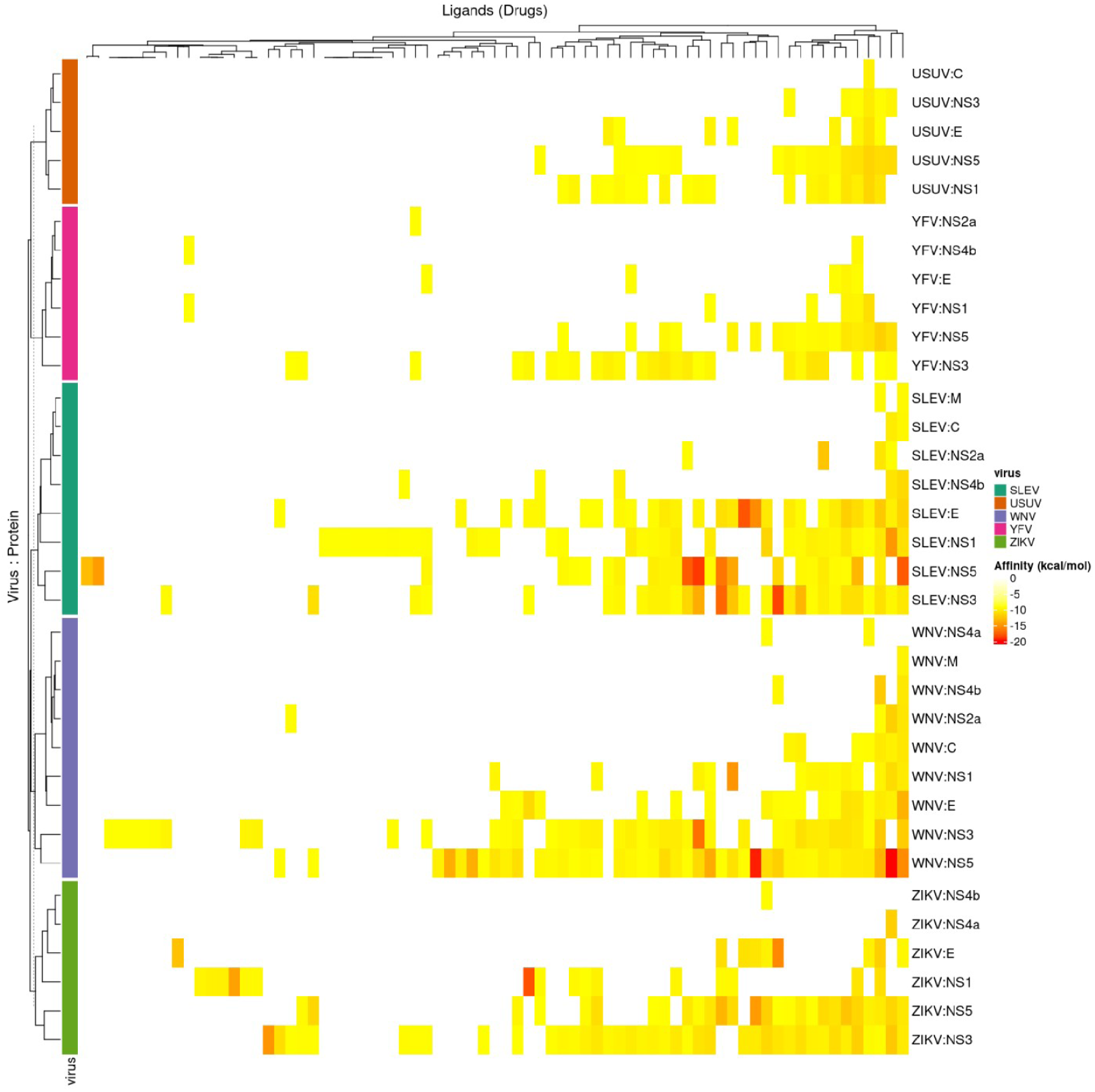
Hierarchical clustering of predicted binding affinities (ΔG, kcal/mol) across the flaviviral proteome. The heatmap displays clustering between viral proteins (rows) and all screened compounds (columns) for the five analyzed flaviviruses: USUV, YFV, SLEV, WNV, and ZIKV. Affinities are shown on a yellow-to-dark-red scale, with darker shades indicating stronger predicted binding (ΔG ≤ –15 kcal/mol). Dendrograms summarize similarity relationships among compounds and among virus–protein combinations, outlining shared pharmacodynamic signatures and cross-reactivity patterns (compound identifiers omitted for clarity).

In Figure 7, compound clusters correspond to multi-target scaffolds such as Suramin, Ivermectin, Quercetin, and Andrographolide, whose co-clustering across phylogenetically distinct proteins highlights conserved recognition modes. These patterns are further supported by complementary dimensionality-reduction analyses, including UMAP and LDA (Supplementary Material, file 1, figures S7 and S8).

Regarding the affinity range, two major protein clusters emerge. The upper branch (enriched in USUV and YFV proteins) shows predominantly moderate affinities, indicating a narrower ligand-recognition profile. The lower branch (containing SLEV, WNV, and ZIKV proteins) displays a dense band of high-affinity interactions, particularly for NS3, NS5, and E, reflecting the physicochemical adaptability of these enzymatic and surface-glycoprotein interfaces. Substructure-specific trends also can be identified: SLEV and WNV (both members of the Japanese encephalitis serocomplex) share strong co-affinity signatures in NS3 and NS5, whereas ZIKV displays a distinct low-affinity block centered on NS4A/NS4B, suggesting divergence in replication-complex architecture.

Functionally, the data show that hydrophobic scaffolds and bulky polyanionic compounds tend to act on Capsid, Envelope, and Membrane proteins, consistent with interference in membrane-dependent entry or egress. In contrast, NS3 and NS5 form a stable cluster across most viruses (except YFV), reflecting ligands that preferentially engage their enzymatic active sites and show limited affinity for structural proteins. NS1 clusters near the structural group, likely reflecting flavonoids as structural-axis inhibitors. This is consistent with NS1’s dual role as a replication cofactor (non-enzymatic) and as a membrane-associated glycoprotein, whose lipid-exposed surfaces resemble those of E and prM/M.

Meanwhile, NS4A, NS4B, NS2A, and NS2B cluster as small transmembrane proteins with hydrophobic binding surfaces that preferentially recruit non-polar ligands such as porphyrins and polycyclic scaffolds. As a result, most hits against these proteins show poor selectivity and behave as non-specific aggregators. Effective inhibition may therefore require alternative strategies (e.g., induced degradation or indirect disruption) or acceptance of a “niclosamide-like” profile — broad, low-specificity activity that can still be therapeutically viable.

Flavonoids and nucleoside analogs also display strong affinity for E and NS5, aligning with divergent interaction clusters. These generally polar to moderately lipophilic scaffolds preferentially engage surface-exposed structural regions and the NS5 polymerase. In contrast, highly non-polar compounds (including temoporfin, chlorin E6, phthalocyanines, and other tetrapyrrolic derivatives) bind strongly to NS3 and the transmembrane proteins (NS2A, NS2B, NS4A, NS4B) but show minimal activity toward E and NS5. This separation indicates that no single compound, in its current form, is likely to cover all target classes; instead, distinct chemotypes specialize in different subsets of viral proteins.

In summary, the results outline a framework supporting a complementary multi-drug strategy for emerging flaviviruses. A rational approach would pair a compound from the polar cluster (e.g., myricetin or sofosbuvir) with one from the non-polar cluster (e.g., temoporfin or niclosamide), leveraging their distinct target preferences. Docking simulations identified several molecules with measurable affinity for multiple proteins across all evaluated flaviviral proteomes (ZIKV, YFV, WNV, SLEV, and USUV). Overall, the data point to two avenues for future work: (i) assessing individual compounds that engage coherent groups of viral targets, and (ii) testing combinations that collectively cover diverse protein classes. Experimental validation will be essential to determine the pharmacological feasibility of these strategies.

### Pan-Viral Multi-Target Compounds (Broad-Spectrum Monotherapy)

A single drug capable of inhibiting multiple stages of the flaviviral cycle across diverse species would represent the most straightforward therapeutic option. Although none of the screened molecules fully meet this criterion, our workflow identified several compounds that approach it. These candidates are readily visible in the bar plot (Supplementary Material, file 1, figure S9), with the most notable examples including:

#### Polyphenolic flavonoids

Molecules such as myricetin, quercetin, and luteolin show strong, recurrent binding to both NS5 polymerase and the E glycoprotein across multiple flaviviruses, suggesting dual interference with genome replication and entry/fusion. These compounds already exhibit documented antiviral activity in vitro (including against ZIKV and DENV through conserved mechanisms) and possess favorable pharmacological profiles: generally low toxicity, oral bioavailability, and well-tolerated metabolism^39, 82^. Their modular chemistry makes them highly amenable to optimization, where small substitutions can enhance potency or stability. Overall, flavonoids remain strong prototype scaffolds for broad-spectrum, multi-target antiviral development^90^.

#### Nucleoside analogs

Sofosbuvir, an approved HCV drug with a well-defined safety profile, primarily targets NS5 but may also accommodate secondary pockets in NS3, potentially affecting both RNA synthesis and helicase activity^77^. Ribavirin, though less potent and dose-limited by cytotoxicity^91^, also emerged on our screen as a ligand of conserved catalytic regions in NS5 and NS3. While ribavirin is unlikely to succeed as monotherapy, its broad tissue distribution and immunomodulatory properties may be valuable in combination regimens. Overall, sofosbuvir shows limited promise as a standalone flaviviral inhibitor, while analogs or combination therapies could enhance its antiviral range^92^.

#### Porphyrinic protease inhibitors

Temoporfin shows strong *in vivo* activity against ZIKV and disrupts NS2B–NS3 complex formation across multiple flaviviruses^88^. However, as emphasized, its systemic clinical use is hampered by extreme hydrophobicity. Therefore, if delivery improvements are achieved, temoporfin or analogs could function as protease-focused monotherapy (preventing polyprotein processing). Even so, protease-only monotherapy is vulnerable to rapid resistance, given the high mutation rate of flaviviral RNA genomes and the likelihood of NS3 escape variants, in addition to possible off-target effects on host proteases. For these reasons, potent NS3 inhibitors like temoporfin, or optimized analogs, are best positioned as components of combination therapy rather than standalone agents.

#### Polyanionic derivatives

Molecules such as ATA and suramin can interfere with several virus–host interactions, including capsid–RNA binding and engagement of viral proteins with cellular cofactors^83, 93^. Suramin is an approved antitrypanosomal drug that inhibits dengue and other flaviviruses by binding the E glycoprotein and potentially interfering with polymerase activity. In our screening, ATA appeared as a representative scaffold of this class but was removed during ADME filtering, so any positive signals should be interpreted with caution. Although polyanions show potential as broad multi-target inhibitors, their clinical utility is limited by toxicity, poor specificity, and delivery challenges; substantial refinement would be required for any viable monotherapy.

Collectively, flavonoids and sofosbuvir stand out by combining broad target coverage with favorable ligand efficiency. In practice, a truly pan-flaviviral agent could greatly simplify outbreak management. Although these candidates still require optimization and experimental validation, our computational pipeline effectively highlights the molecules with the strongest translational potential.

### Therapeutic Combination Strategies (Multi-Target Attack)

Building on the multi-target patterns described above, combination therapy is a logical extension. Our findings support several rational pairings:

#### Protease Inhibitor + Polymerase Inhibitor

This combination could provide a broader resistance barrier than either agent alone. By blocking proteolytic processing upstream (e.g., temoporfin) and RNA-chain elongation downstream (e.g., sofosbuvir), the virus encounters two mechanistically independent checkpoints during replication. This layered pressure is particularly relevant for flaviviruses, whose NS3–NS5 interplay is highly coordinated and whose mutational escape routes are constrained by essential enzymatic functions. Docking data indicate that sofosbuvir and ribavirin bind in proximate catalytic regions of NS5 and NS3, suggesting synergistic pressure on the core replicative machinery. Moreover, adopting an HCV-inspired dual-inhibitor strategy leverages a clinically validated paradigm within a similarly polyprotein-driven replication cycle.

#### Envelope (E) Inhibitor + Protease/Polymerase Inhibitor

This combination could simultaneously prevent entry and egress (via E inhibition) while blocking intracellular genome replication (via NS3/NS5). A practical example is pairing a flavonoid E-binder (e.g., myricetin) with sofosbuvir or with a protease inhibitor such as niclosamide. A myricetin–sofosbuvir regimen would offer a fully oral, low-cost combination with a potentially favorable safety profile. Our heatmaps reinforce the rationale: E and NS5 fall into distinct target clusters (Figure 7), supporting a division of labor between two dedicated agents. This computational complementarity aligns with experimental evidence of compounds exerting broad antiviral effects, possibly through E protein destabilization in its pre-fusion state ^94, 95^.

#### Ligands targeting NS1 or membrane proteins (NS4A/4B)

Although NS1, NS4A, and NS4B lack well-validated inhibitors compared to classical enzymatic targets, there is a strong mechanistic rationale for incorporating them into therapeutic strategies. Circulating NS1 contributes to immune evasion and endothelial dysfunction^96^, so blocking its oligomerization or secretion could attenuate pathogenesis and complement a replication inhibitor.

For NS4A/NS4B, a combination such as niclosamide + sofosbuvir is a plausible option, especially given that comparable pairings have shown promise in other viral systems (e.g., simeprevir synergizing with remdesivir against SARS-CoV-2)^97^. In the flavivirus context, niclosamide has a well-established safety range and may function as an adjuvant that enhances the penetration or efficacy of co-administered drugs within replication compartments. Accordingly, a three-pronged regimen combining (i) an entry inhibitor (E), (ii) a replicase inhibitor (NS3/NS5), and (iii) a membrane microenvironment modulator (targeting NS4A/4B and related functions) could provide a higher and more resilient level of viral suppression.

#### Inhibitors of viral protein-protein interactions

Another combination strategy is to pair catalytic inhibitors with molecules that disrupt essential non-enzymatic interactions. For instance, a recent study identified a small molecule that interferes with the NS3–NS5 interaction, a contact required for proper assembly of the flaviviral replication complex^98^. One can then envision a triple regimen comprising an NS5 inhibitor (e.g., sofosbuvir), an NS3 inhibitor (e.g., temoporfin or a derivative), and a third agent that destabilizes the NS3–NS5 interface. Incorporating such orthogonal (allosteric) mechanisms creates a high barrier to resistance, as the likelihood of the virus acquiring simultaneous compensatory mutations across multiple targets (E, NS3, NS5, etc.) becomes extremely small under the selective pressure of three coordinated drugs.

The development of antiviral cocktails must account for pharmacokinetics and the risk of cumulative adverse effects. Even so, several compounds highlighted in our screening are either approved drugs or naturally present in the human diet, suggesting a generally favorable safety margin. It is also worth noting that polytherapy broadens antiviral coverage, an advantage in regions where coinfections are possible or the etiological agent is initially uncertain. In such cases, a combination regimen ensures that at least one drug is effective against each pathogen. For instance, niclosamide exhibits activity against the Chikungunya virus (an alphavirus)^99^, so a niclosamide + sofosbuvir combination could potentially address scenarios of Zika and Chikungunya coinfection during concomitant outbreaks.

### Rationale for the Multi-Target Attack and Supporting Evidence

The clinical literature in antivirals reinforces our data towards a modular approach: in HIV, for example, a protease inhibitor + two reverse transcriptase inhibitors are used, ensuring that maturation and genomic replication “modules” are blocked^100^. In the case of flaviviruses, our proposed modules are: (1) viral entry/exit (E, C, prM, NS1), (2) RNA replication (NS5 polymerase, NS3 helicase), and (3) polyprotein processing and replicative complex assembly (NS3 protease, NS2B, NS4A, NS4B). Based on the workflow presented, we could identify an optimal representative ligand, respectively.

Additional support for combination therapy comes from the dual-binding behavior of certain compounds. For instance, flavonoids targeting both NS5 and E indicate that these proteins share pockets accommodating similar chemistries, revealing potential overlapping structural vulnerabilities. Additionally, we noted that ribavirin and sofosbuvir interact with conserved catalytic motifs of NS5 and NS3, which suggests that nucleoside analogs can inhibit multiple viral enzymes (ribavirin itself acts by causing mutagenesis in the polymerase and general energy deficit by depleting intracellular GTP)^91^.

It is important to emphasize that using multiple drugs from the start of treatment can prevent the emergence of resistant strains - a common scourge when using antiviral monotherapy. For ZIKV and other flaviviruses, resistance risk is lower than in HIV due to their acute, short-duration infections; however, single-drug pressure can still select for mutants during prolonged outbreaks or in immunocompromised hosts. Resistance to protease inhibitors has already been observed in DENV under monotherapy in cell culture^101^.

## TABLE OF MATERIALS

A Summary of Software, Databases, and computational resources used in the Multi-Target Docking and ADME Analysis Pipeline is provided below. [Place **Table of Materials** here]

## DISCUSSION

The primary contribution of this work is the implementation of a transparent, reusable, and auditable computational software workflow that integrates pocket detection, docking, ADME descriptors, ligand-efficiency metrics, and multivariate prioritization. The workflow provides a structured and reproducible pocket-oriented virtual screening and chemoinformatics framework for comparative profiling of compounds across multiple flaviviral targets. It is intended to support hypothesis generation, retrospective benchmarking, and downstream experimental prioritization rather than to serve as stand-alone proof of antiviral efficacy.

The workflow is particularly suited for projects involving multiple molecular targets and large compound libraries, as it enables simultaneous evaluation across homologous systems and supports meaningful comparisons that reveal both expected and unexpected binding patterns (as exemplified by structural and non-structural protein coverage). Additionally, this workflow offers several key advantages for computational drug discovery. First, it restricts computational tools to three languages (R, Python, and Bash), ensuring accessibility and ease of customization while leveraging free and reputable software. This design makes the workflow suitable for beginners in bioinformatics by providing a ready-to-run framework that minimizes complex computational functions and reduces the need for multiple iterative runs. Second, the integrated multivariate analysis capabilities (e.g., PCA, t-SNE, UMAP, and LDA) enable sophisticated exploration of chemical space and target–compound relationships, which is essential for broader drug-discovery projects104. Third, the modular architecture allows expert users to modify and extend functionality, such as incorporating additional descriptors or customizing the Structural Complexity Index (SCI) calculation, thereby improving workflow capabilities without extensive reprogramming.

Another notable methodological feature of the workflow is the use of concavity analysis to explore binding patterns beyond classical catalytic-site interactions. This enables bias-mitigated investigations of polymer-surface adaptations, including allosteric interference, perturbation of the protein–membrane interface, and polarity-driven contributions that may be essential for pharmacological optimization and could inform more sophisticated analyses employing Molecular Dynamics105, 106. This approach improves reliability and hypothesis refinement by systematically exploring multiple binding sites (pockets) per target and capturing an extensive range of predicted interactions, thereby providing comprehensive insight into compound–target relationships that may guide therapeutic strategies against flaviviruses. Through this methodological implementation, the study proposes candidate compounds for single or combination therapies, informed by the potential for synergistic interactions.

From a public health standpoint, the concept of an adaptable therapeutic cocktail against flaviviruses—modeled after combined antiretroviral therapy—holds strong translational potential. Accordingly, the multi-target framework proposed here offers at least three key advantages for multidisciplinary research groups responding to recent outbreaks: robustness, by maintaining efficacy despite viral mutation; flexibility, by informing therapeutic adjustments according to structural homology; and computational efficiency, through automated large-scale screening of compounds across multiple viral targets. This enables a stepwise path toward experimental validation: (i) biochemical assays to measure enzyme inhibition (e.g., NS3 protease, NS5 polymerase), (ii) antiviral assays in cell culture to assess reductions in viral replication, and (iii) cytotoxicity profiling to evaluate compound safety. Concurrently, lead compounds identified through this automated screening can be further evaluated using additional LBVS and SBVS techniques^107^. In this study, several promising candidates were also identified as suitable for extended investigation in animal models, particularly when supported by established preclinical evidence (e.g., sofosbuvir^77^, temoporfin^108^, and niclosamide^109^).

The workflow addresses computational constraints by minimizing redundant calculations, automating data aggregation, and enabling efficient matrix operations for large-scale analyses. In addition, automated file detection and processing eliminate most manual intervention and reduce the risk of errors through integrated critical-step checks. These features make the workflow particularly valuable for screening campaigns where computational resources are limited or where rapid iteration is needed to evaluate multiple compound libraries against evolving target panels. Considering this, the workflow can also be extended beyond flavivirus research to other multi-target drug discovery applications, including antimicrobial resistance, cancer therapy, and neurodegenerative diseases, as its modular design facilitates adaptation to diverse target families, compound collections, and pharmacological requirements.

Despite these advantages, several limitations should be acknowledged. The workflow relies on molecular docking predictions, which, although computationally efficient, may not fully capture dynamic protein–ligand interactions or solvation effects under physiological conditions110, 111. The ADME descriptors are derived from static molecular structures and, therefore, may not reflect metabolic transformations or tissue-specific distribution111. Moreover, the multivariate analyses assume linear or locally linear relationships in chemical space, which may overlook more complex interaction patterns in highly diverse compound libraries. Notwithstanding, the workflow offers a rational and computationally efficient foundation for pre-clinical studies, illustrating how integrated computational approaches can accelerate broad-spectrum therapeutic design and guide experimental validation toward the most promising candidate compounds even in resource-aware research environments.

### Troubleshooting

Common technical issues encountered during computational screening and data processing are summarized in Table 5. Proper dataset curation and environment configuration are critical to ensure reproducibility and stability of the workflow. [Place **Table 5** here]

**Table 5.** Most common technical issues, potential causes, and recommended solutions encountered in this computational workflow.

The workflow scripts, example directory structure, and documentation are available in the associated repository (see Methods, step 1.8). For software-oriented dissemination, users and developers are encouraged to archive an exact release (e.g., via Zenodo), include a minimal runnable example dataset, document software versions and dependencies, and provide expected output files to facilitate reproducibility and independent reuse.

## Supporting information

Workflow

Violin

LLE vs PKd

Heatmap

2D map

H. Clustering

## DATA AND CODE AVAILABILITY

## ACKNOWLEDGMENTS

This work was supported by the National Council for Scientific and Technological Development (CNPq) under Grant Nos. 442831/2023-4 and 309271/2022-3, and by the Coordination for the Improvement of Higher Education Personnel—Brazil (CAPES—Financing Code 001) in accordance with Ordinance 206 of 04/09/2018.

We dedicate this work, in memoriam, to Prof. Dr. Carlos Francisco Sampaio Bonafé, whose remarkable scientific legacy continues to inspire us.

