## Supplementary figures and images for "An Open-Source Reproducible Workflow for Pocket-Oriented Virtual Screening and ADME-Integrated Chemoinformatics: A Multi-Target Flavivirus Case Study"

### 2D map

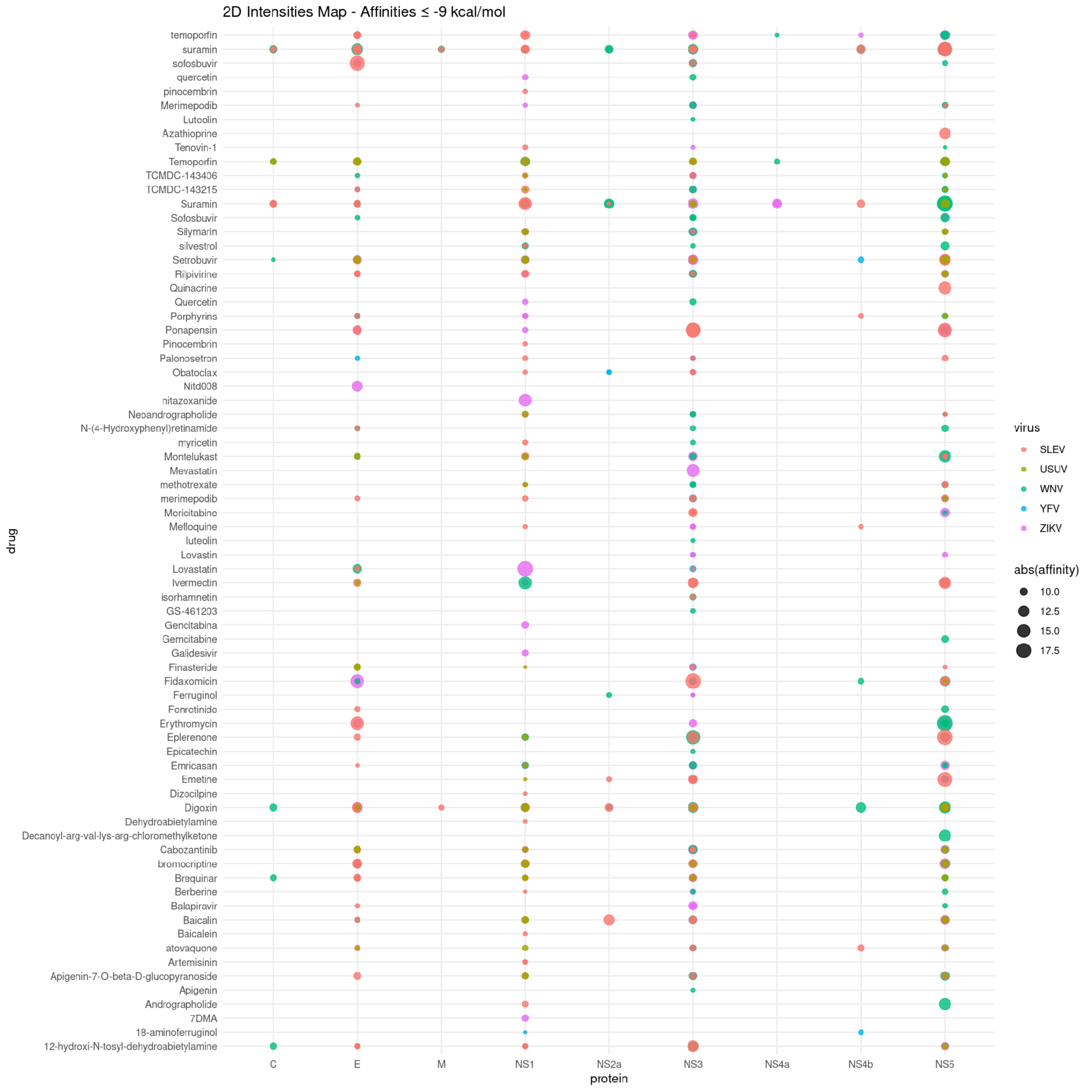

### H. Clustering

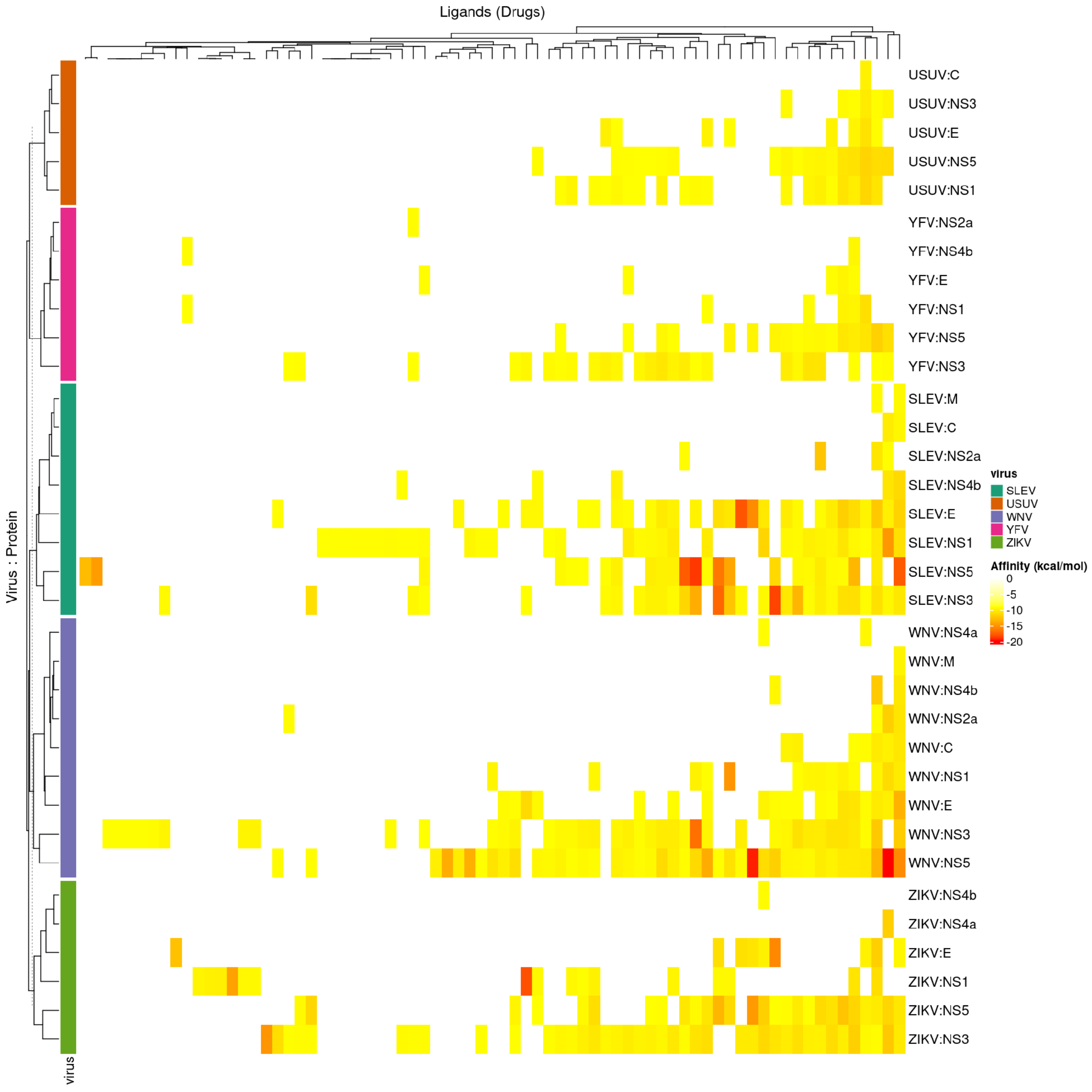

### Heatmap

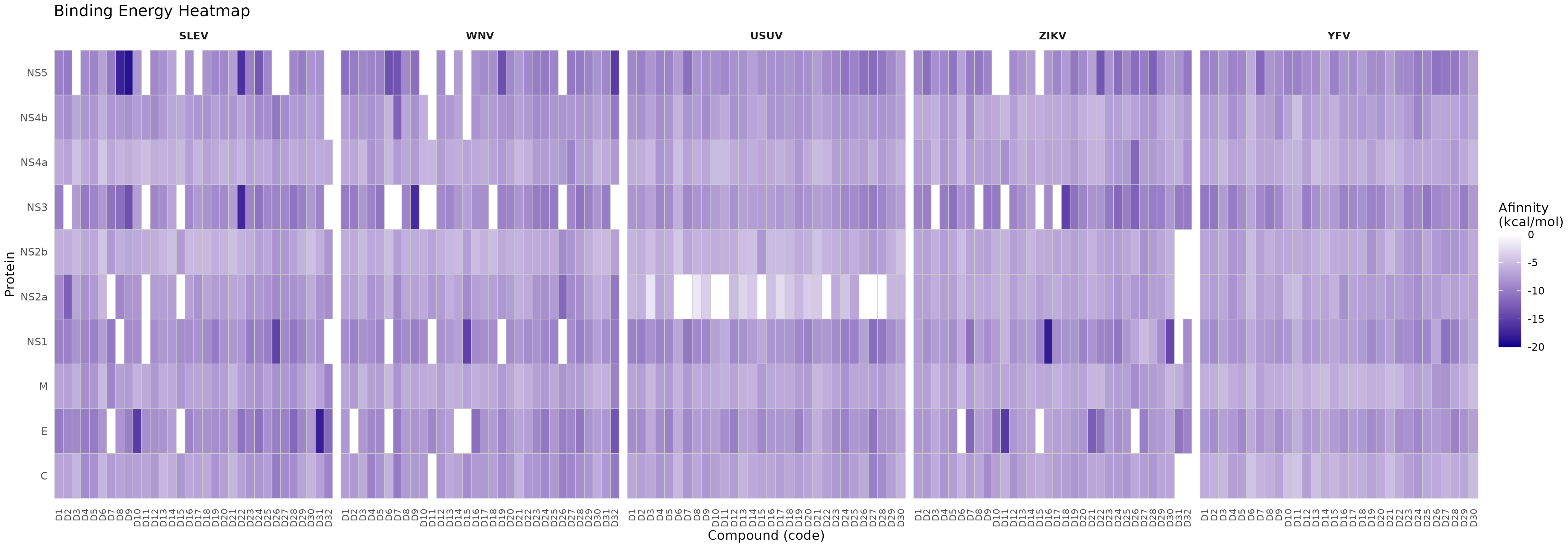

### LLE vs PKd

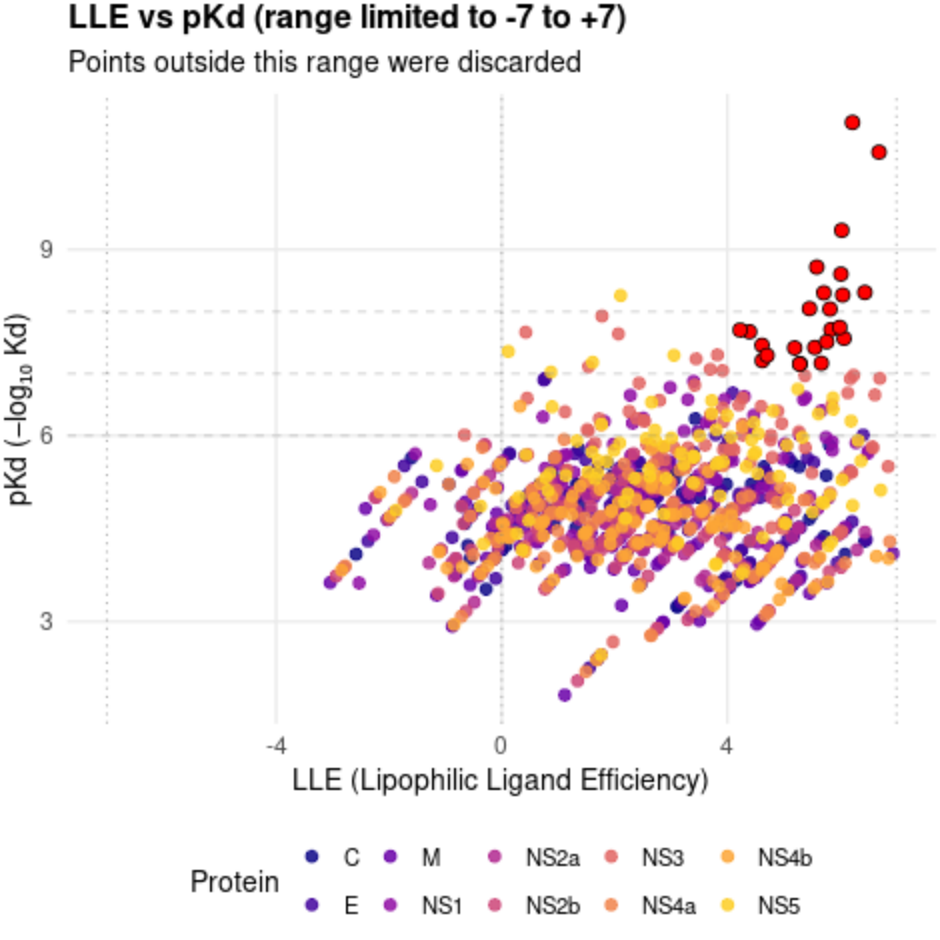

### NS5

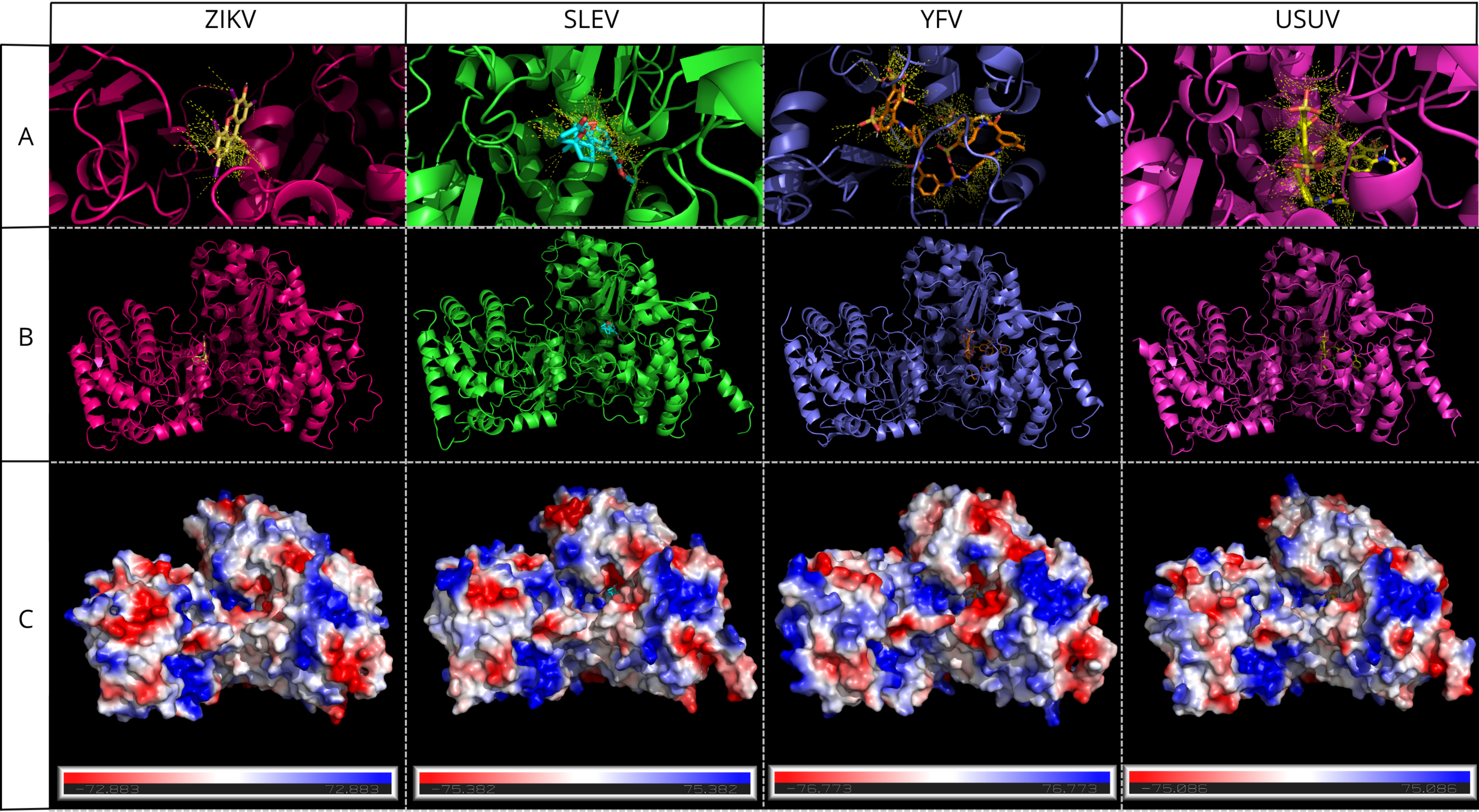

### Violin

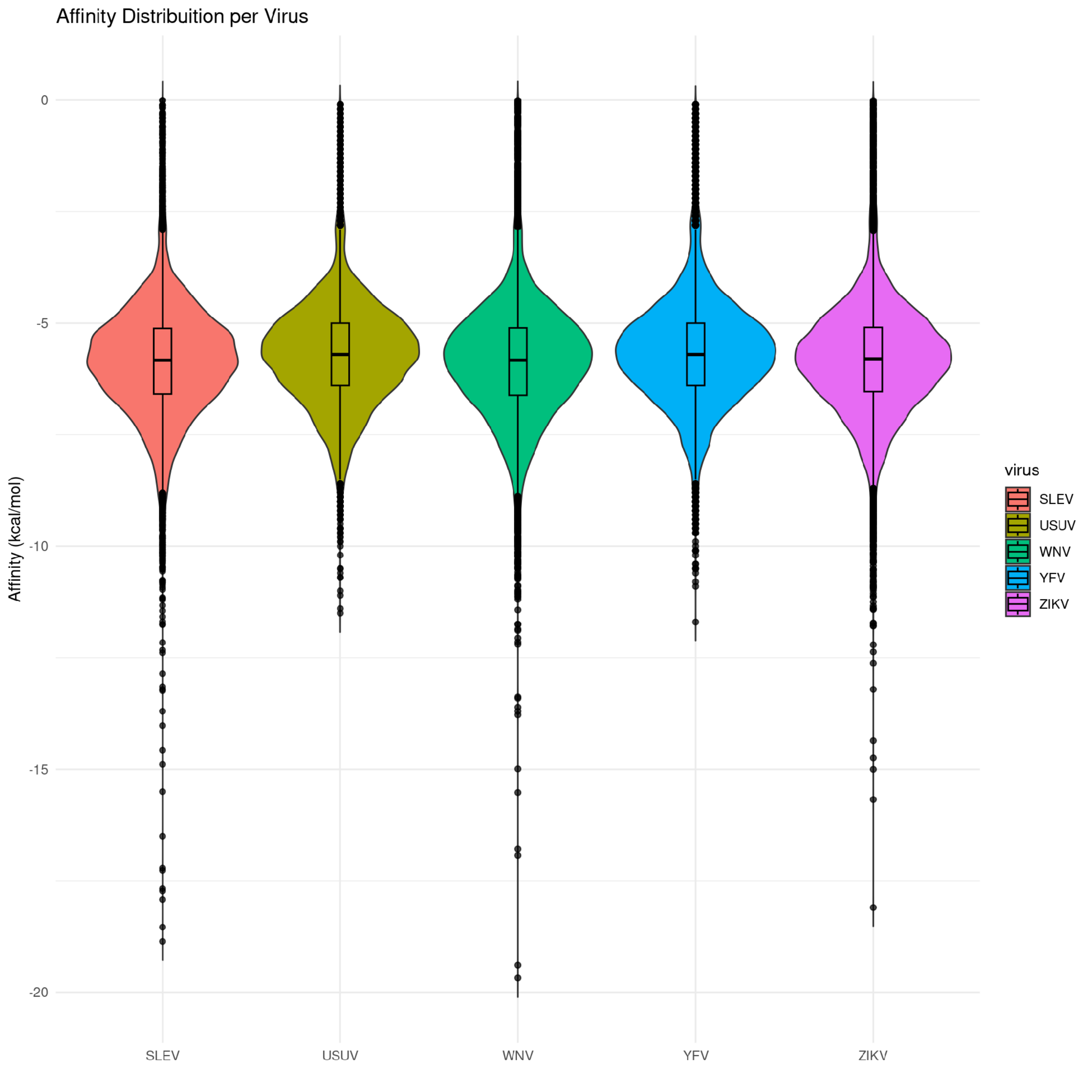

### Workflow

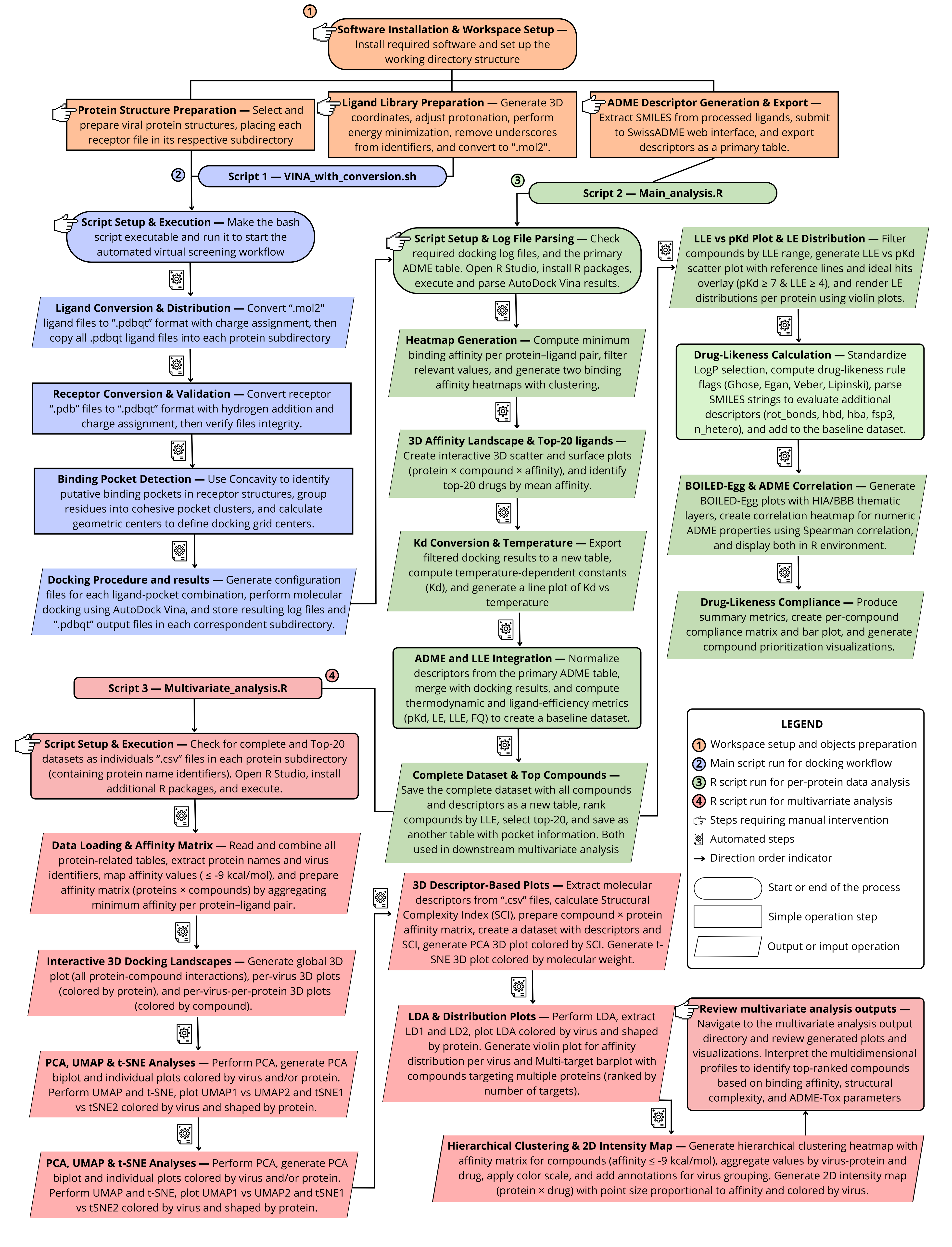
